# Serine Stabilizes SLC7A11 and Enhances Cystine Influx to Protect Against Acute Pancreatitis

**DOI:** 10.64898/2026.05.02.722375

**Authors:** Yan Huang, Fei Fu, Lingli Deng, Yiqin Wang, Jiawang Li, Jiaxian Zhang, Jinxi Yang, Yijing Long, Manjiangcuo Wang, Chenxia Han, Lihui Deng, Pengfei Li, Haiyang Chen, Jiyang Dong, Xianghui Fu, Qing Xia, Dan Du

**Author notes:** Correspondence to (D. Du), Frontiers Science Center for Disease-related Molecular Network, West China Hospital, Sichuan University, Chengdu 610041, China. Correspondence to (Q. Xia), Department of Integrated Traditional Chinese and Western Medicine, West China Hospital, Sichuan University, Chengdu 610041, China. Correspondence to (X. Fu), Department of Biotherapy, Center for Diabetes and Metabolism Research, State Key Laboratory of Biotherapy and Cancer Center, West China Hospital, Sichuan University, Chengdu, 610041, China. Correspondence to (J. Dong), Department of Electronic Science, National Institute for Data Science in Health and Medicine, Xiamen University, Xiamen 361005, China. These authors made equal contributions to this work (Yan Huang, Fei Fu, Lingli Deng).

## Abstract

Lethal sterile inflammatory diseases are linked to amino acid metabolism, but the role of serine remains unclear. Here, we show that dysregulated serine metabolism and reduced plasma serine levels correlate with disease severity of acute pancreatitis (AP) in patients and mouse models. Elevating serine levels via exogenous serine supplementation or pancreatic phosphoglycerate dehydrogenase (PHGDH) overexpression mitigates pancreatic injury, whereas a serine deprivation diet or pancreatic PHGDH knockdown exacerbates AP. Serine prevents cell death and oxidative stress in pancreatic acinar cells, human induced pluripotent stem cells-derived pancreatic organoids and mouse pancreatic tissue. Serine enhances cysteine and glutathione biosynthesis primarily by promoting solute carrier family 7 member 11 (SLC7A11)-dependent cystine uptake rather than by serving as a direct substrate. Mechanistically, the E3 ubiquitin ligase NEDD4 mediates ubiquitination and degradation of SLC7A11, whereas serine binds to NEDD4 and thereby inhibits SLC7A11 degradation. Similarly to serine, pharmacological inhibition of NEDD4 alleviates lipid peroxidation and pancreatic injury. These findings identify serine as a critical signaling regulator of SLC7A11 stability and oxidative stress, and provides a new therapeutic strategy for AP and associated sterile inflammatory disorders.

**Highlights:** Acute pancreatitis (AP) is linked to abnormal serine metabolism and serine depletion.

Serine prevents cell death in AP acinar cells, human pancreatic organoids and mice.

Serine promotes SLC7A11-dependent cystine uptake and glutathione levels in acinar cells.

Serine reduces NEDD4-mediated ubiquitination of SLC7A11.

**In brief:** Serine protects against cell death and pancreatic injury in acute pancreatitis by stabilizing SLC7A11 through disruption of NEDD4-mediated ubiquitination in acinar cells.

## INTRODUCTION

Lethal sterile inflammatory conditions including severe trauma, acute liver failure, acute pancreatitis (AP), and ischemia-reperfusion injury are life-threatening, and non-infectious disorders characterized by rapid onset and a paucity of effective pharmacotherapies ^1,2^. In these settings, the innate immune system is overactivated by damage-associated molecular patterns (DAMPs) released from injured or necrotic cells, leading to systemic inflammatory response syndrome (SIRS), multiple organ dysfunction syndrome (MODS), and even death ^3^. Emerging evidence has established metabolic stress and dysregulation as key pathological drivers in inflammatory cascades, with amino acid metabolism demonstrating a robust correlation with disease severity particularly in AP ^4,5^. As a prototypical sterile inflammatory disorder of the exocrine pancreas, AP is initiated by acinar cell intrinsic insults, including oxidative stress, premature intracellular trypsinogen activation, mitochondrial dysfunction, and diverse forms of cell death; these events collectively drive DAMPs release, local pancreatic inflammation, and subsequent SIRS ^6–10^. Cysteine and glutathione (GSH) metabolism plays a critical role in maintaining redox homeostasis in pancreatic acinar cells ^11^. However, the involvement of additional amino acid metabolites in modulating this pathway remains incompletely defined. Therefore, systematic identification of disease-relevant metabolic checkpoints in AP holds significant translational potential for developing mechanism-informed therapeutic interventions.

Serine is increasingly recognized as a conditionally essential amino acid obtained from the diet or synthesized *de novo* via the phosphorylated pathway ^12^. It serves as a central metabolic hub, providing carbon and nitrogen for the biosynthesis of GSH, sphingolipids, and nucleotides, as well as cysteine through condensation with homocysteine ^13^. Beyond its metabolic roles, serine functions as a signaling molecule, allosterically activating pyruvate kinase M2 (PKM2) to regulate glycolysis and redox metabolism, with PKM2 activity markedly reduced under serine deprivation ^14^. Clinically, serine deficiency is associated with liver injury and ischemia-reperfusion injury ^15,16^, whereas serine supplementation restores redox balance and alleviates lipotoxicity in animal models ^17^. Although serine supports the biosynthesis of cysteine and sphingolipids ^11^, both critical for maintaining redox homeostasis and restraining sterile inflammatory responses in AP, its therapeutic potential and the molecular mechanisms underlying its modulation of AP pathogenesis remain incompletely defined. Elucidating these regulatory functions is essential for developing targeted, metabolism-based therapies.

Solute carrier family 7 member 11 (SLC7A11) is a cystine/glutamate antiporter essential for maintaining cellular redox homeostasis in diverse cell types, including pancreatic acinar cells ^18–20^. By importing cystine, which is reduced to cysteine, SLC7A11 provides the rate-limiting substrate for GSH synthesis, a key ROS scavenger ^21^. Dysregulation of SLC7A11 has been linked to liver injury, ischemia-reperfusion injury and inflammatory conditions ^22–25^. In severe AP animal models, reduced SLC7A11 expression contributes to lipid peroxide accumulation and pancreatic injury ^20,26–30^. However, the post-transcriptional or post-translational mechanisms controlling its abundance remain largely unknown. Ubiquitin-mediated degradation via E3 ligases, such as SOCS2 and TRIM7, regulates SLC7A11 stability in cancer ^31,32^, but whether similar mechanisms operate in AP is unclear. Notably, impaired cystine uptake can induce *de novo* serine biosynthesis ^33^, suggesting a potential interplay between serine metabolism and SLC7A11 function. Understanding how serine regulates SLC7A11 expression and cystine transport is therefore critical for maintaining redox homeostasis in acinar cells and preventing disease progression.

Here, we demonstrated that serine deficiency is closely related to severe AP in both patients and mouse models, and supplementary serine alleviates lipid peroxidation and pathological damage in AP, whereas serine deficiency exacerbates injury. We also report a previously unrecognized mechanism in acinar cells: serine directly stabilizes SLC7A11 protein by disrupting neural precursor cell expressed, developmentally down-regulated 4 (NEDD4) mediated ubiquitination. This stabilization significantly enhances cystine influx, restores intracellular cysteine and GSH levels, and suppresses lipid peroxidation and inflammation in pancreatic acinar cells, organoids, or murine model levels. Our findings unexpectedly identify serine as a post-translational regulator of transporter stability that couples amino acid metabolism to proteostasis, thereby sustaining cystine import and redox balance. This work not only expands the therapeutic repertoire of serine but also unveils the NEDD4-SLC7A11-cystine influx axis as a critical determinant of acinar cell survival and a promising target for AP and related inflammatory disorder intervention.

## RESULTS

### Reduced plasma serine concentration is significantly associated with increased disease severity in patients with acute pancreatitis

Metabolic features that are consistently dysregulated in both human patients and preclinical animal models are critically implicated in the pathogenesis of AP. To identify the disease-specific metabolic pathway exhibiting the greatest perturbation in AP, we first constructed individual concentration perturbation profiles (ICPP) using a personalized perturbation framework adapted from transcriptomic analysis ^34^ (**Fig. 1a**). It integrates previously acquired targeted metabolomics data from AP patients ^35^ with newly generated targeted metabolomics datasets from two timecorse murine models including cerulein (CER)-induced mild edematous AP and arginine (ARG)-induced severe necrotizing AP. The top 15 disease-specific pathways in AP patients (**Fig. 1b and Supplementary Tab. 1**), the top 12 in CER-induced AP mice (**Fig. 1c and Supplementary Tab. 2**), and the top 16 in ARG-induced AP mice (**Supplementary Fig. 1a and Supplementary Tab. 3**) were therefore identified. Glycine, serine, and threonine metabolism was consistently identified as a key disease-specific pathway in both patients and murine models of AP, ranking among the top 3 perturbed pathways in the severe disease stage.

**Fig. 1:**
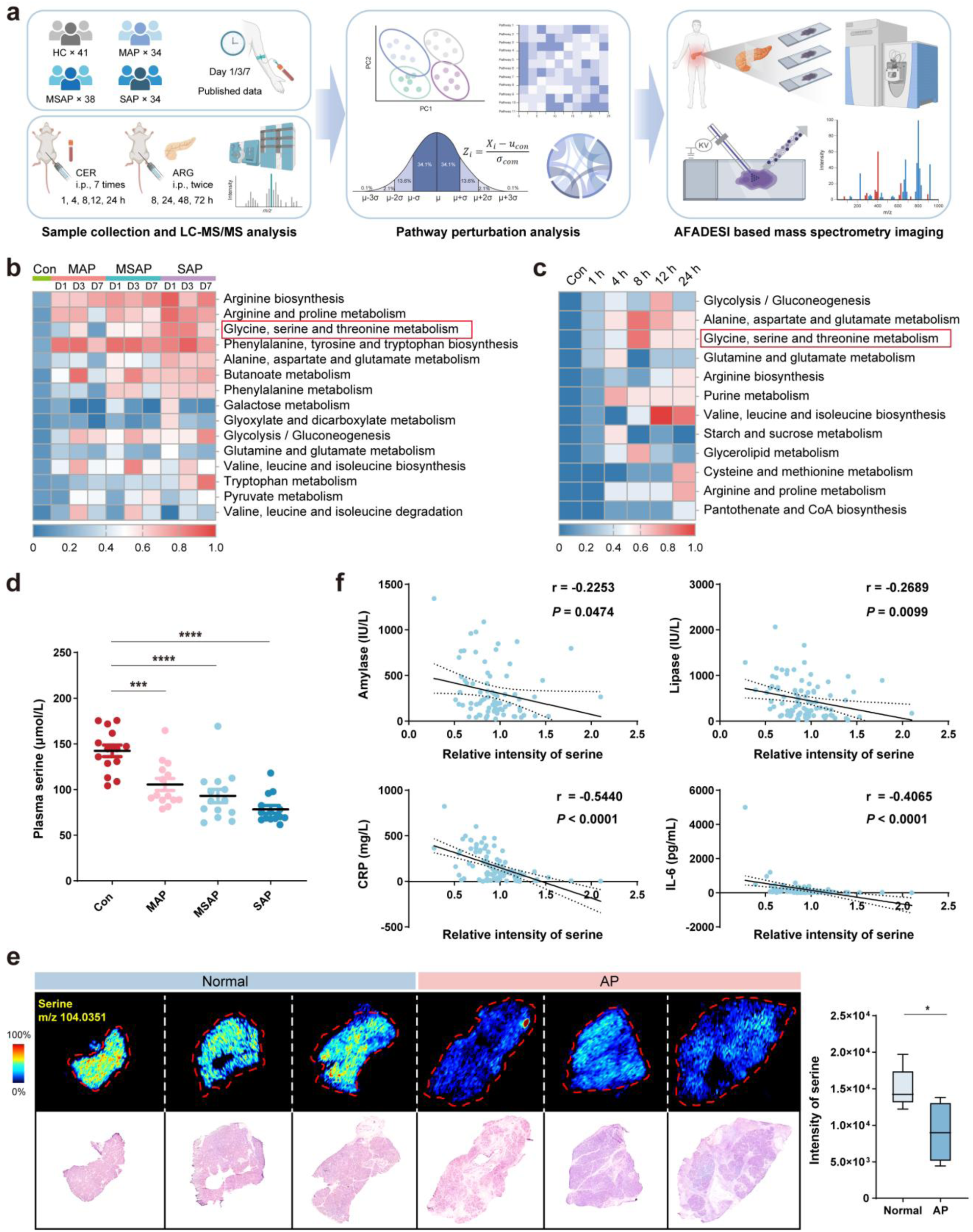
Reduced plasma serine concentration is significantly associated with increased disease severity in patients with acute pancreatitis. a, Schematic overview of the experimental workflow, including targeted metabolomics performed in AP patients and mouse models, personalized metabolite perturbation calculation, and MSI analysis. **b,** Top 15 disease-specific metabolic pathways interrupted in plasma of AP patients versus healthy controls (HCs), identified by ICPP-based pathway perturbation analysis (adjusted *P* < 0.01). Targeted metabolomics were analyzed using 321 plasma samples obtained from 106 patients [34 mild AP (MAP), 38 moderately severe AP (MSAP), 34 SAP], who were sampled at three time points [day 1 (D1), day 3 (D3), and day 7 (D7) after admission], together with 41 HCs. **c,** Top 12 disease-specific metabolic pathways interrupted in serum of CER-induced AP mice, identified by ICPP-based pathway perturbation analysis (adjusted *P* < 0.01). Targeted metabolomics were analyzed using serum samples from CER-induced AP mice at five timepoints (1 h, 4 h, 8 h, 12 h, 24 h after model establishment) and control mice (n = 7). **d,** Absolute quantitation of serine concentrations in plasma from AP patients and HCs by LC-MS/MS (n = 14). **e,** Representative MSIs showing serine distribution in resected human pancreas and quantitative analysis of serine intensities from AP patients and paracancerous controls. Right panel: Quantitative analysis of serine signal intensity was performed in three separate 5×5 pixel regions for each tissue (n = 4–5). **f,** Pearson correlations of plasma serine intensity with amylase, lipase, CRP and IL-6 levels (n = 78–95). Values shown are mean ± SEM. * *P* < 0.05, *** *P* < 0.001, and **** *P* < 0.0001.

Further evaluation of pathway metabolite perturbations across AP severity stages revealed that in comparison to other metabolites, serine changed the most significantly (Z-score =-2.31) and was present at the lowest level in severe AP (SAP) (**Supplementary Fig. 1b–d**). Absolute quantification of plasma serine by liquid chromatography-tandem mass spectrometry (LC-MS/MS) in AP patients further validated this finding (**Fig. 1d**). Mass spectrometry imaging (MSI) of necrotic pancreatic tissue from SAP patients and adjacent normal tissue from pancreatic cancer patients clearly demonstrated significantly reduced serine concentrations in AP (**Fig. 1e**). These observations establish that serine reduction is a typical feature of AP. Correlation analysis revealed that serine levels were significantly and inversely correlated with clinical diagnostic markers amylase and lipase, as well as prognostic biomarkers C-reactive protein (CRP) and interleukin 6 (IL-6), relative to other metabolites in the glycine, serine and threonine pathway (**Fig. 1f and Supplementary Fig. 1e,f**). Collectively, these findings indicate that dysregulation of serine metabolism, particularly the reduction in serine concentration is closely associated with AP progression.

### Exogenous and endogenous modulation of serine availability regulates the severity of acute pancreatitis

Serine is derived from dietary intake or synthesized *de novo* via phosphoglycerate dehydrogenase (PHGDH) (**Fig. 2a**). First, to evaluate the therapeutic effect of exogenous serine, we administered serine in a CER-induced mild AP model (**Fig. 2b**). Serine administration significantly increased serine levels in both serum and pancreas (**Fig. 2c**), reduced CER-induced elevation in serum amylase and lipase as well as pancreatic myeloperoxidase (MPO) activities (**Fig. 2d-g**), and alleviated pancreatic oedema, and mild acinar cell necrosis (**Fig. 2h**). Serine also restored the ultrastructural integrity of acinar cells by reducing mitochondrial swelling and endoplasmic reticulum distension (**Fig. 2i**). Consistently, in ARG-induced severe necrotizing AP (**Supplementary Fig. 2a**), serine supplementation similarly elevated serine levels in both serum and pancreas (**Supplementary Fig. 2b**), reduced serum amylase and lipase activities as well as pancreatic MPO activity (**Supplementary Fig. 2c**), and ameliorated pancreatic inflammatory cells infiltration, and extensive acinar cell necrosis (**Supplementary Fig. 2d**). These protective effects were accompanied by improved mitochondrial morphology and reduced endoplasmic reticulum stress (**Supplementary Fig. 2e**). In comparison, mice were subjected to a serine-and glycine-deficient diet (SGD) for 4 weeks prior to CER-AP induction (**Supplementary Fig. 3a**). This SGD dietary intervention significantly reduced serum serine levels (**Supplementary Fig. 3b**), confirming effective depletion of dietary serine. Compared to mice maintained on a normal diet, SGD-fed mice exhibited increased lipase activities and elevated pancreatic MPO activity (**Supplementary Fig. 3c**), and exacerbated inflammatory infiltration and acinar necrosis following AP induction (**Supplementary Fig. 3d**). Together, these results indicate that increasing serine availability alleviates AP severity, whereas serine deficiency exacerbates AP.

**Fig. 2:**
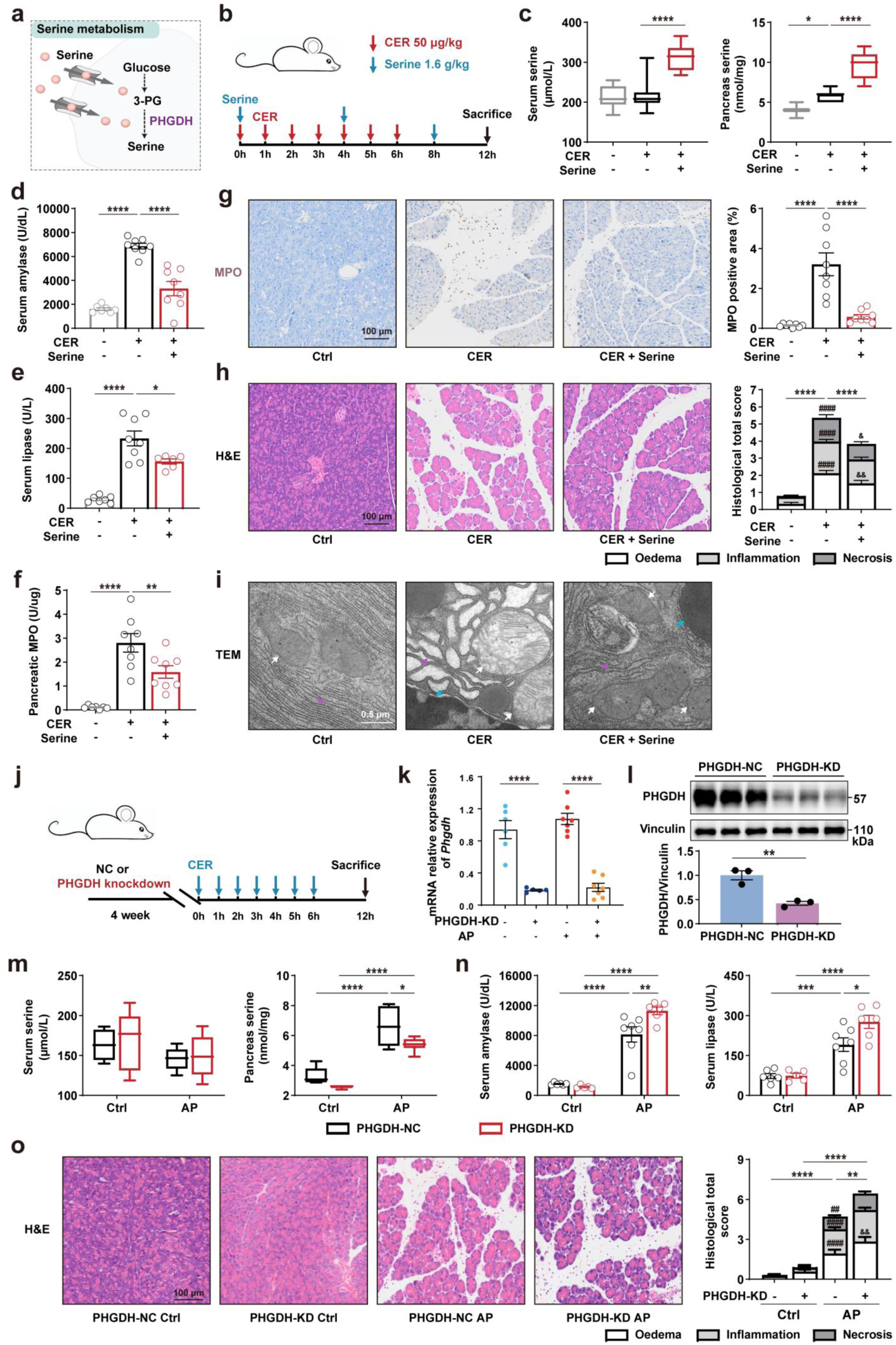
Exogenous and endogenous intervention of serine concentration regulates the severity of acute pancreatitis. a, Schematic illustration of serine uptake via *de novo* synthesis and membrane transport. **b,** Schematic of CER-induced AP modeling and serine administration protocol. **c,** Serum and pancreatic serine concentrations (n = 7–8). **d-f,** Serum (**d**) amylase and (**e**) lipase activities, along with (**f**) pancreatic MPO activity (n = 6–8). **g,** Representative IHC images and quantification of pancreatic MPO expression (n = 8). **h,** Representative pancreatic histological images (H&E staining) and quantitative histopathological scoring (right panel) for oedema, inflammation, and necrosis subscores (n = 6–8). Differences in total scores are denoted by *. For subscores, # represents significance compared with the control group, and & represents significance compared with the AP group. **i,** Transmission electron microscopy (TEM) images showing ultrastructural changes in pancreatic tissue (White arrows, mitochondria; purple arrows, endoplasmic reticulum; green arrows, nucleus). **j,** Schematic illustration of pancreas-specific PHGDH knockdown via retrograde pancreatic duct injection, followed by CER-induced AP modeling. **k,l,** Quantification of PHGDH expression in pancreatic tissue: (**k**) mRNA levels by RT-qPCR (n = 5–7) and (**l**) protein levels by western blot (with Vinculin as loading control, n = 3). **m,** Serine concentrations in serum and pancreatic tissue, measured by LC-MS/MS in PHGDH-KD versus PHGDH-NC (negative control) groups with or without CER induction (n = 5–7). **n,** Serum amylase and lipase activities (n = 5–7). **o,** Representative pancreatic histological images (H&E staining) and quantitative histopathological scoring (right panel) for oedema, inflammation, and necrosis subscores (n = 5–7). Differences in total scores are denoted by *. For subscores, # represents significance compared with the PHGDH-NC Ctrl group and & represents significance compared with the PHGDH-NC AP group. Values shown are mean ± SEM. * *P* < 0.05, ** *P* < 0.01, *** *P* < 0.001, **** *P* < 0.001.

To assess the role of endogenous serine synthesis, we modulated PHGDH expression in the pancreas. Pancreas-specific PHGDH knockdown (PHGDH-KD) using adeno-associated virus (AAV)-mediated short hairpin RNA (**Fig. 2j**) significantly reduced PHGDH mRNA and protein levels (**Fig. 2k,l**). In CER-AP, PHGDH-KD mice exhibited lower pancreatic serine versus PHGDH-NC mice (**Fig. 2m**), higher serum amylase and lipase activities (**Fig. 2n**), more severe pancreatic oedema (**Fig. 2o**). Conversely, pancreas-specific PHGDH overexpression (PHGDH-OE) using AAV delivered via retrograde pancreatic duct injection (**Supplementary Fig. 4a**). Successful AAV transduction was confirmed by enhanced green fluorescent protein (EGFP) expression localized to the pancreatic parenchyma (**Supplementary Fig. 4b**) and significant upregulation of *Phgdh* mRNA in the PHGDH-OE group relative to the negative control group (PHGDH-NC) (**Supplementary Fig. 4c**). PHGDH-OE increased the serine level in the pancreas (**Supplementary Fig. 4d**) and attenuated CER-induced elevations in serum amylase and lipase as well as pancreatic trypsin and MPO activities (**Supplementary Fig. 4e,f**). Compared to the PHGDH-NC group, PHGDH-OE mice displayed attenuated pancreatic oedema, inflammatory cell infiltration, and acinar necrosis during AP **(Supplementary Fig. 4g)**. Ultrastructural analysis further showed improved mitochondrial morphology and reduced ER dilation in acinar cells (**Supplementary Fig. 4h**). Collectively, these findings demonstrate that both exogenous supplementation and endogenous enhancement of serine biosynthesis increase serine availability and mitigate disease severity in AP.

### Serine protects against oxidative stress and cell death in primary acinar cells and human iPSC-derived pancreatic organoids

To clarify the primary cell types of serine’s action in AP, we performed single-cell RNA sequencing (scRNA-seq) on pancreatic tissue from CER-AP mice with or without serine supplementation (schematic in **Supplementary Fig. 5a**). Unsupervised clustering of 29 initial clusters (**Supplementary Fig. 5b,c**), followed by cell annotation (12 major population) and sub-clustering of the acinar compartment, 3 transcriptionally distinct acinar subpopulations were resolved (**Fig. 3a–d**). It includes *Pdia2* high functional acinar cells, *Krt8* high acinar-to-ductal metaplasia (ADM)-like acinar cells, and *Ccr2* high inflammatory acinar cells (**Fig. 3e**). CER-AP drastically shifted this distribution, expanding the *Ccr2*-high inflammatory subset while depleting functional and ADM-like cells. Serine treatment largely reversed these changes, restoring the proportions of functional and ADM-like acinar cells and reducing the inflammatory subpopulation to near-control levels (**Fig. 3e and Supplementary Fig. 5d**). Within the acinar compartment, serine further counteracted CER-induced downregulation of antioxidant genes (*Gpx1, Gpx4, Sod1, Sod2, Cat, Nfe2l2* and *Hmox1*) (**Supplementary Fig. 5e**), indicating potential antioxidant modulation in acinar cells.

**Fig. 3:**
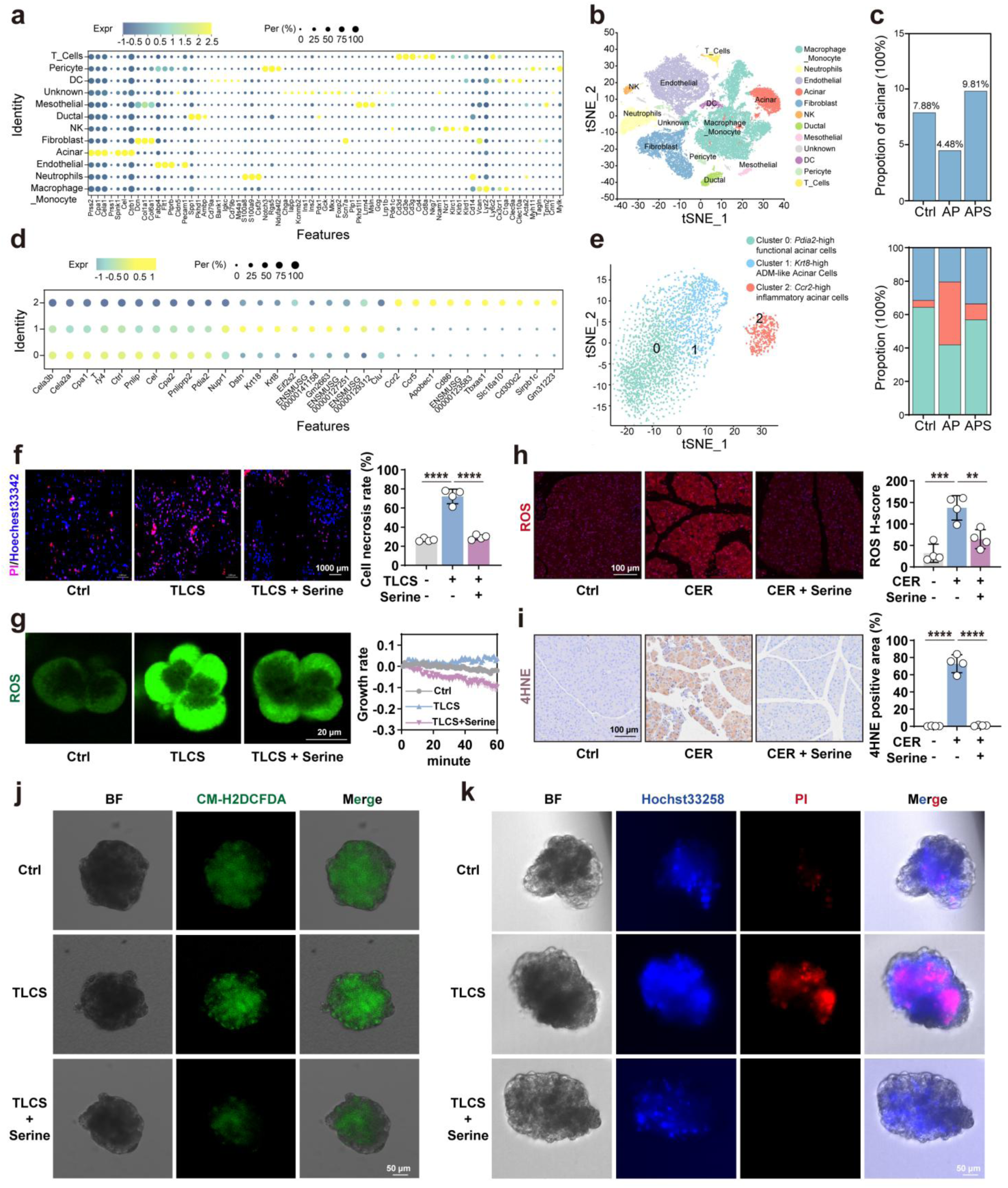
Serine protects against oxidative stress and cell death in PACs and human iPSC-derived pancreatic organoids. a, Bubble plot displaying marker gene expression levels used to annotate cell types in scRNA-seq data from pancreatic tissue. Bubble size indicates the percentage of cells expressing the marker; color intensity represents the average expression level. **b,** t-SNE dimensionality reduction plot of all pancreatic cells, colored by annotated cell types. **c,** The proportion chart of acinar cells in the three groups. **d,** Dot plot showing differential expression of unique differentially expressed markers across the three identified acinar cell subclusters. **e,** tSNE plot highlighting three acinar cell clusters (left panel), and the proportional composition of the three acinar cell subclusters in Ctrl, AP, and APS groups (right panel). **f,** Representative fluorescence images of pancreatic acinar cell necrosis with PI/hoechst staining in TLCS-treated PACs with or without serine pretreatment, and quantitative necrosis rate (% PI-positive cells) (n = 4). **g,** Representative real-time fluorescence traces of ROS generation (H₂DCFDA probe) in living PACs during TLCS stimulation with or without serine pretreatment. **h,** Representative fluorescence images and quantification of pancreatic ROS expression in CER-AP mice with or without serine treatment (n = 4). **i,** Representative IHC images and quantification of pancreatic 4-HNE expression in CER-AP mice with or without serine treatment (n = 4). **j,** Representative fluorescence images of ROS levels (H₂DCFDA probe) in TLCS-induced human iPSC-derived pancreatic organoids with or without serine pretreatment. **k,** Representative fluorescence images of cell death in TLCS-induced pancreatic organoids, stained with PI (dead cells) and hoechst (all nuclei), with or without serine pretreatment. Values shown are mean ± SEM. ** *P* < 0.01, \*\*\**P* < 0.001, and **** *P* < 0.0001.

To evaluate the cytoprotective effects of serine on acinar cells, we first performed *ex vivo* experiments using isolated mouse primary acinar cells (PACs). Propidium iodide (PI) /hoechst staining demonstrated that serine reduced taurolithocholic acid 3-sulfate (TLCS)-induced necrosis in a dose-dependent manner, with optimal protection at approximately 100 mM (**Fig. 3f and Supplementary Fig. 6a**). Mechanistically, serine enhanced superoxide dismutase (SOD) and glutathione peroxidase (GPX) activities to eliminate ROS accumulation (**Fig. 3g and Supplementary Fig. 6b,c**) and mitigated AP-associated ATP depletion induced by various toxins (**Supplementary Fig. 6d**). In pancreatic tissue, serine administration reduced CER-induced ROS accumulation and 4-HNE levels (**Fig. 3h,i)**. Furthermore, PHGDH-KD mice exhibited higher pancreatic ROS and 4-HNE levels compared to PHGDH-NC mice, whereas PHGDH-OE mice showed lower ROS and 4-HNE levels than PHGDH-NC controls (**Supplementary Fig. 6e-h)**.

For clinical relevance, we generated human iPSC-derived pancreatic organoids (**Supplementary Fig. 7a**). Morphological progression from iPSC (day 0) to day-11 progenitors and day-15 organoids was confirmed (**Supplementary Fig. 7b**). Immunofluorescence revealed decreasing expression of pancreatic and duodenal homeobox 1 (PDX1, a marker of all pancreatic progenitor cells) and NK6 homeobox 1 (NKX6.1, a marker of trunk progenitor cells), as well as increasing expression of amylase (AMY, a marker of acinar cells), from day 12 to 15, with amylase localizing to peripheral regions (**Supplementary Fig. 7c**). RT-qPCR verified elevated acinar and ductal marker gene expressions by day 15 (**Supplementary Fig. 7d**), confirming successful differentiation into acinar and ductal lineages. In TLCS or H_2_O_2_-injured organoids, serine treatment improved the morphological characteristics (**Supplementary Fig. 7e**). Moreover, serine reduced ROS production, particularly in peripheral acinar-like regions, decreased cell death, and increased the level of ATP (**Fig. 3j,k and Supplementary Fig. 7f**). Collectively, serine supplementation preserves acinar cell identity, reduces inflammatory acinar populations, and directly protects acinar cells from oxidative stress and cell death.

### Serine enhances cysteine and GSH production primarily through stabilization of SLC7A11 and increasing cystine influx

To elucidate how serine protects acinar cells, we performed targeted metabolomics on pancreatic tissues from CER-AP mice (**Supplementary Tab. 4**). Principal component analysis (PCA) revealed distinct clustering among groups and KEGG enrichment identified “cysteine and methionine metabolism” and “glutathione metabolism” as one of the most significantly affected pathways (**Supplementary Fig. 8a**), consistent with our scRNA-seq data linking serine to oxidative stress protection. In PACs, TLCS-induced reduction of cysteine and GSH levels, which were reversed by serine treatment (**Fig. 4a**), and *in situ* GSH imaging further confirmed the restoration by serine (**Fig. 4b**). We next investigated whether serine is directly involved in cysteine and GSH metabolic pathways (**Fig. 4c**). Isotope-labeled metabolic flux experiments revealed minimal contribution from *de novo* synthesis or extracellular serine: tracing with [¹³C_6_]-glucose revealed cysteine labeling (<1%), and [¹³C_3_]-serine tracing yielded similarly low labeling (<5%) (**Fig. 4d**), indicating that endogenous synthesis is not the primary source. In contrast, [¹³C_6_]-cystine tracing demonstrated rapid uptake and reduction to cysteine, achieving nearly 40% labeling within 5 min, a process significantly enhanced by serine treatment (**Fig. 4e**). These results establish serine-enhanced cystine uptake as the predominant source of cysteine in acinar cells.

**Fig. 4:**
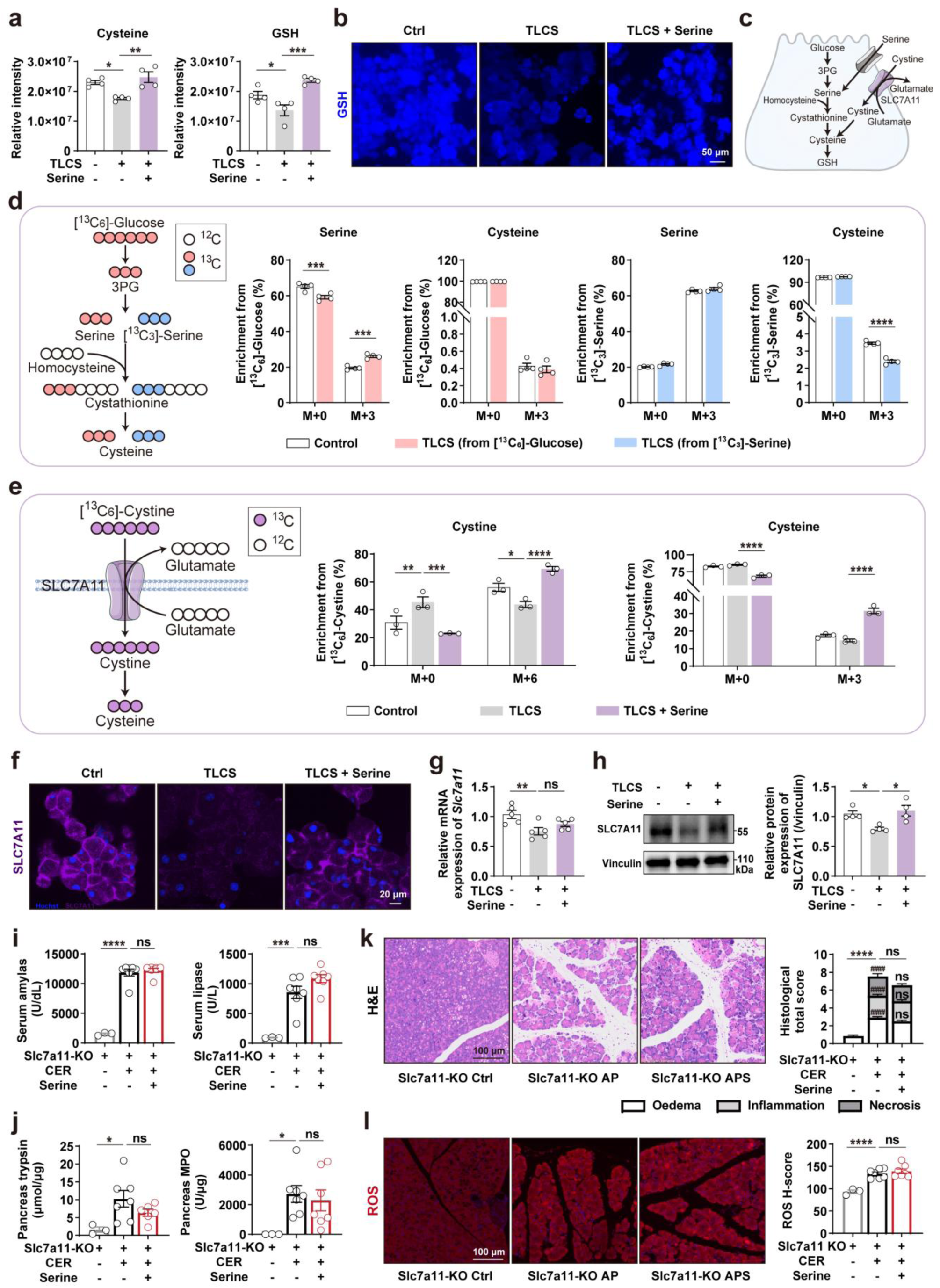
Serine enhances cysteine and GSH production primarily through stabilization of SLC7A11 and increasing cystine influx. a, Relative quantitation of intracellular cysteine and GSH levels in PACs following TLCS induction, with or without serine pretreatment (n = 4), using a derivatization-based LC-MS/MS assay. **b,** Representative immunofluorescence images of intracellular GSH in PACs following TLCS induction, with or without serine. **c,** Schematic diagram illustrating the pathways of cysteine generation and metabolism in PACs. Cysteine is derived from *de novo* synthesis through glucose and serine metabolism, as well as from SLC7A11-mediated transport. **d,** Isotope labeling ratios of serine and cysteine in PACs cultured with [¹³C₆]-glucose (2 h) or [¹³C₃]-serine (30 min) tracers (n = 4). Data represent fractional labeling. **e,** Isotope labeling ratios of cystine and cysteine in PACs labeled with [¹³C₆]-cystine (5 min, n = 3). Data represent fractional labeling. **f,** Immunofluorescence images of SLC7A11 in PACs following TLCS induction and serine pretreatment. **g,** Relative *Slc7a11* mRNA expressions in PACs (n = 5). **h,** Western blot analysis and quantification of SLC7A11 protein levels in PACs (n = 4). **i,** Serum amylase and lipase activities (n = 3–7). **j,** Pancreatic trypsin and MPO activities (n = 3–7). **k,** Representative pancreatic histological images (H&E staining) and quantitative histopathological scoring (right panel) for oedema, inflammation, and necrosis (subscores and total score) (n = 3–7). Differences in total scores are denoted by *. For subscores, # represents significance between control and AP groups, and & represents significance between APS and AP groups. **l,** Representative fluorescence images and quantification of pancreatic ROS levels (n = 3-7). Values shown are mean ± SEM. * *P* < 0.05, ** *P* < 0.01, *** *P* < 0.001, and **** *P* < 0.0001. ns meaning *P* > 0.05.

Cystine transport is predominantly mediated by SLC7A11. Immunofluorescence revealed high SLC7A11 expression on the plasma membrane of PACs, with additional cytoplasmic localization (**Fig. 4f**). TLCS stimulation markedly reduced SLC7A11 protein levels but not mRNA levels, which were restored by serine treatment (**Fig. 4g,h**). Consistent results were observed in pancreatic tissue from ARG-AP models (**Supplementary Fig. 8b–d**), supporting a post-translational mechanism of SLC7A11 stabilization regulated by serine. Genetic ablation of Slc7a11 abolished serine’s protective effects against CER-induced pancreatitis, as evidenced by no significant changes in serum amylase and lipase, pancreatic tissue trypsin and MPO activities, as well as pathological damage, and ROS and 4-HNE accumulation (**Fig. 4i–l and Supplementary Fig. 8e-g**), demonstrating that SLC7A11 is essential for this serine-mediated effect. These results together clearly demonstrate that serine exerts its antioxidant function by stabilizing SLC7A11 protein, thereby promoting cystine uptake and subsequent cysteine and GSH synthesis.

### Serine directly binds the E3 ubiquitin ligase NEDD4 and reduces NEDD4-mediated SLC7A11 ubiquitination

To investigate whether serine regulates SLC7A11 degradation through the ubiquitin-proteasome or lysosomal pathway (**Fig. 5a**), we performed proteomic analysis, which implicated ubiquitin-related pathways (**Supplementary Fig. 9a and Supplementary Tab. 5**). Functional assays confirmed that SLC7A11 is degraded primarily via the ubiquitin-proteasome system, as its levels were stabilized by the proteasome inhibitor MG-132 but not by a lysosomal inhibitor chloroquine (CQ) (**Fig. 5b,c**). Consistently, TLCS-induced increased SLC7A11 ubiquitination levels in both PACs and 266-6 cells, an effect attenuated by serine treatment (**Fig. 5d,e**). Ubiquitination involves a cascade of E1, E2 and E3 enzymes, with E3 ligases specifying substrates. To identify the exact enzyme regulate this process, we performed immunoprecipitation-mass spectrometry (IP-MS) with an anti-SLC7A11 antibody, revealing the E3 ubiquitin ligase NEDD4 as a candidate (**Fig. 5f,g and Supplementary Tab. 6**). Co-IP experiments confirmed that NEDD4 interacts with SLC7A11, enhanced by TLCS induction in PACs and in 266-6 cells by MG132 treatment (**Fig. 5h,i**). NEDD4 overexpression reduced SLC7A11 protein levels and increased its ubiquitination (**Fig. 5j,k**), establishing NEDD4 as a novel E3 ligase for SLC7A11.

**Fig. 5:**
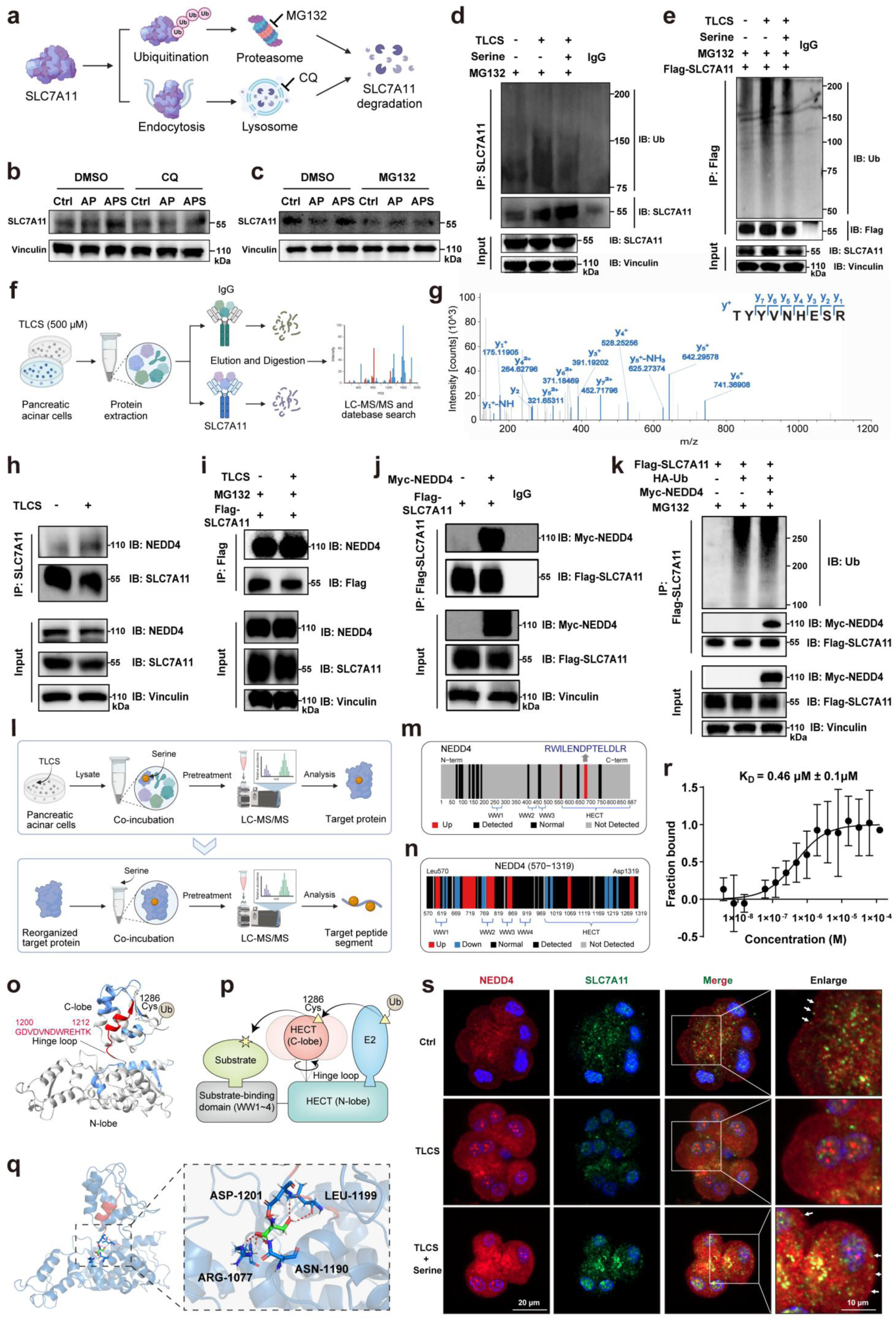
Serine directly binds the E3 ubiquitin ligase NEDD4 and reduces NEDD4-mediated SLC7A11 ubiquitination. a, Diagram of the protein degradation process of SLC7A11. **b,c,** WB analysis of SLC7A11 protein stability in PACs treated with (**b**) the lysosomal inhibitor CQ (50 μM, 30 min) or (**c**) the proteasome inhibitor MG132 (10 μM, 30 min) with or without serine pretreatment. Vinculin served as a loading control. **d,e,** Immunoprecipitation (IP) followed by WB detection of SLC7A11 ubiquitination in TLCS-treated (**d**) PACs and (**e**) 266-6 cells, with or without serine pretreatment and MG132 (10 μM, 4 h) to accumulate ubiquitinated forms. IP was performed with anti-SLC7A11 antibody; Ubiquitin was detected with anti-Ub. Input lysates are shown below. **f,** Experimental workflow for identifying SLC7A11-interacting E3 ubiquitin ligases via IP-MS in TLCS-treated PACs. **g,** Representative tandem mass spectrometry and peptide sequences of NEDD4 identified as an SLC7A11-interacting protein in TLCS-treated PACs. **h,** Endogenous Co-IP confirming interaction between NEDD4 and SLC7A11 in PACs (IP with anti-SLC7A11). **i,** Exogenous Co-IP in 266-6 cells overexpressing Flag-SLC7A11, treated with MG132. IP with anti-Flag; interaction detected by blotting for the reciprocal tag. **j,** WB analysis of SLC7A11 and NEDD4 protein levels in 266-6 cells overexpressing Flag-SLC7A11 transfected with empty vector (control) or Myc-NEDD4 overexpression plasmid. Vinculin as a loading control. **k,** Detection of SLC7A11 ubiquitination in 266-6 cells overexpressing Flag-SLC7A11 transfected with the indicated plasmids (control vector, Myc-NEDD4, or HA-Ub) and treated with MG132. IP with anti-Flag; ubiquitin detected with anti-HA antibody. **l,** Schematic illustration of LiP-MS workflow to identify serine-binding proteins and conformational changes in PACs lysates and recombinant protein. **m,** Conformational alterations of NEDD4 induced by serine treatment in PACs lysates. Significantly altered LiP peptides (*P* < 0.05, and FC ≥ 2 or FC ≤0.5) are mapped onto the complete sequence of mouse NEDD4, presented in QR code format. **n,** Conformational alterations in human recombinant NEDD4 protein induced by serine treatment. Significantly altered LiP peptides (*P* < 0.05, and FC ≥ 1.5 or FC ≤ 0.67) are mapped onto the human NEDD4 sequence. **o,** Significantly differentially accessible peptides mapped onto the three-dimensional structure of the NEDD4 HECT domain (PDB: 2XBF). **p,** The domain architecture of NEDD4, showing the WW domains and catalytic HECT domain, along with the corresponding ubiquitin transfer process. **q,** Molecular docking model of serine binding to NEDD4 (HECT domain), with detailed protein-ligand interaction diagram (The red dotted lines indicate hydrogen bonds). **r,** MST analysis showing the interaction of serine with recombinant NEDD4. **s,** Representative immunofluorescence images of PACs co-stained for SLC7A11 (green) and NEDD4 (red), showing subcellular colocalization. Nuclei counterstained with DAPI.

Next, to determine how serine modulates NEDD4-mediated SLC7A11 ubiquitination, we employed limited proteolysis-mass spectrometry (LiP-MS) in PACs (**Fig. 5l**). Screening revealed 226 out of 1,534 proteins exhibiting significant conformation alterations (*P* < 0.05, and fold change, FC ≥ 2 or FC ≤ 0.5; **Supplementary Fig. 9b–e and Supplementary Tab. 7**). NEDD4 was among the candidates, with conformational changes localized to its HECT domain, while SLC7A11 was absent (**Fig. 5m and Supplementary Tab. 7**). These results suggest that serine directly binds NEDD4 and modulates its conformation. In order to increase peptide coverage, we used recombinant human NEDD4 protein, LiP-MS confirmed serine-induced structural alterations, with 121 of 546 peptides showing significant changes (*P* < 0.05, and FC ≥ 1.5 or FC ≤ 0.67, **Supplementary Fig. 9f,g and Supplementary Tab. 8**). Binding sites were mapped to the WW1-WW3 substrate-recognition domains and the HECT catalytic domain (**Fig. 5n**). A key conformational shift occurred in the flexible hinge loop of the HECT domain, which regulates ubiquitin transfer and catalytic activity (**Fig. 5o,p**). Molecular docking indicated serine forms hydrogen bonds with residues ASP1201, LEU1199, ARG1077, and ASN1190 in this region (**Fig. 5q**). Microscale thermophoresis (MST) and surface plasmon resonance (SPR) experiments confirmed direct binding between serine and NEDD4, with an equilibrium dissociation equilibrium constant (KD) of 0.46 µM and 9.278 µM, respectively (**Fig. 5r and Supplementary Fig. 9h**). Consistent with these findings, serine treatment restored TLCS-induced disruption of NEDD4-SLC7A11 colocalization (**Fig. 5s**). Consequently, serine directly binds the E3 ubiquitin ligase NEDD4, alter its conformation and prevents NEDD4-mediated SLC7A11 ubiquitination and degradation, thereby stabilizing SLC7A11.

#### Pharmacological inhibition of NEDD4 ameliorates oxidative stress and severity in AP

Given that serine exerts its downstream effects through direct binding to NEDD4, we next investigated whether pharmacological inhibition of NEDD4 could recapitulate the protective effects of serine. We treated cells with heclin, a broad HECT domain E3 ligase inhibitor, and assessed its impact on acinar cell injury. PI/hoechst staining revealed that heclin protected PACs from TLCS-induced necrosis in a dose-dependent manner, with optimal protection observed at 2.5 μM (**Supplementary Fig. 10a**). Heclin treatment significantly reduced TLCS-stimulated ROS accumulation (**Supplementary Fig. 10b–d**) and increased intracellular GSH levels (**Supplementary Fig. 10e**), indicating direct cytoprotection through modulation of oxidative stress.

Based on these *in vitro* findings, we evaluated the therapeutic potential of heclin in the CER-induced AP mouse model. Mice received intraperitoneal injections of heclin at 1 mg/kg or 5 mg/kg following AP induction (**Fig. 6a**). Heclin at the dose of 5 mg/kg significantly reduced CER-induced elevation in serum amylase and lipase, and decreased pancreatic MPO activites (**Fig. 6b–d**). Notably, both doses significantly attenuated pathological damage during AP (**Fig. 6e)**. And both doses decreased pancreatic ROS levels and 4-HNE accumulation (**Fig. 6f,g**), confirming antioxidant effects *in vivo*. Taken together, pharmacological inhibition of NEDD4 with heclin ameliorates oxidative stress and severity in AP, suggesting that NEDD4 may be a key target in AP.

**Fig. 6:**
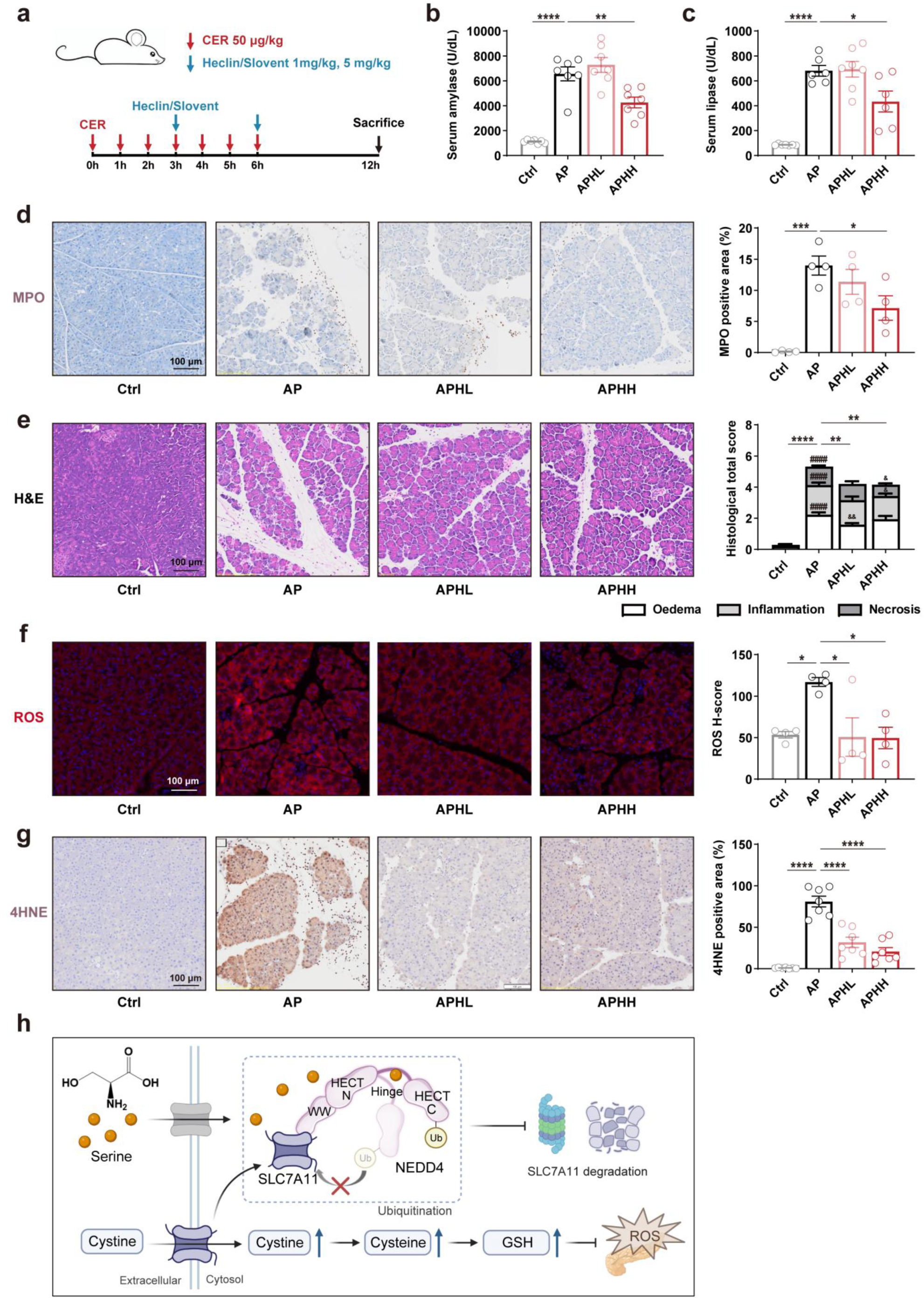
Pharmacological inhibition of NEDD4 with heclin alleviates AP severity in mice. a, Schematic. illustration of CER-induced AP modeling and heclin administration. **b,c,** Serum (**b**) amylase and (**c**) lipase activities (n = 6–7). **d,** Representative IHC images of pancreatic MPO expression (n = 4). **e,** Representative pancreatic histological images (H&E staining) and quantitative histopathological scoring (right panel) for oedema, inflammation, and necrosis (subscores and total score) (n = 6–7). Differences in total scores are denoted by *. For subscores, # represents significance compared with the ctrl group, and & represents significance compared with the AP group. **f,**. Representative fluorescence images of pancreatic ROS levels (n = 4). **g,** Representative IHC images of pancreatic 4-HNE expression (n = 7). **h,** Schematic model illustrating the mechanism of serine action. Values shown are mean ± SEM. * *P* < 0.05, ** *P* < 0.01, *** *P* < 0.001, and **** *P* < 0.0001.

## DISCUSSION

This study reveals a previously unappreciated role for serine as a signaling molecule that directly governs redox homeostasis in AP by modulating the NEDD4-SLC7A11-cystine axis (**Fig. 6h**). Through integrative targeted metabolomics across human and mouse samples, combined with pancreas-specific genetic manipulation and pancreatic organoid models, we demonstrate that serine serves not only as a potential diagnostic biomarker for AP, particularly SAP, but also as a valuable therapeutic supplement for AP. Mechanistically, using *Slc7a11* knockout mice, stable isotope tracing, and molecular interaction analyses in acinar cells, we provide evidence that serine directly binds to the hinge loop of the E3 ubiquitin ligase NEDD4, inducing conformational changes that suppress NEDD4-mediated ubiquitination and degradation of the cystine transporter SLC7A11. This, in turn, stabilizes SLC7A11, enhances cystine uptake, as well as cysteine and GSH synthesis, thereby reducing oxidative stress. Together, these findings establish a direct mechanistic link between serine and SLC7A11 protein homeostasis, uncovering a previously unappreciated axis that governs metabolic reprogramming and cell fate decisions in AP, with broader implications for other lethal sterile inflammatory disease pathophysiology.

### Clinical relevance and translational potential of serine

Emerging evidence indicates profound metabolic disturbances during AP progression ^5,36^, yet a systematic characterization of shared metabolic alterations between human and animal models is lacking. Applying the ICPP algorithm to peripheral metabolomics data from AP patient cohorts of varying severity and corresponding mouse models, we identified glycine, serine and threonine metabolism as a disease-specific pathway, particularly pronounced in SAP. In MAP, this pathway showed transient perturbation—mild perturbation on day 1, peak disruption on day 3, and near normalization by day 7. In contrast, moderately severe (MSAP) and SAP patients exhibited persistent activation across days 1, 3, and 7, with SAP patients failing to fully recover by day 7. These changes are consistent with previous reports of glycine metabolism dysregulation and reduced plasma serine in AP ^37^. Serine emerged as the dominant metabolite within this pathway (Z-score: SAP = −2.31, compared with glycine =-0.72, guanidinoacetate =-2.11, threonine =-1.60, glycerate =-2.16 and betaine =-1.59), correlating most strongly with biochemical markers such as CRP, highlighting its potential as a diagnostic indicator for SAP. At the tissue level, serine concentrations were markedly reduced in resected pancreatic samples from SAP patients and showed a declining trend in SAP animal models, whereas MAP models exhibited a compensatory increase, mirroring prior experimental observations ^38^. Together, these findings suggest that AP onset drives elevated pancreatic serine demand, depleting peripheral levels. In severe AP, enhanced pancreatic serine mobilization and catabolism amplified this depletion, underpinning the pronounced decline in circulating serine observed in SAP.

Although serine supplementation has shown protective effects in diverse inflammatory conditions, including diabetes ^39,40^, intestinal injury ^41^, sepsis ^42^, temporal lobe epilepsy ^43^, white matter demyelination ^44^, and liver fibrosis ^16^, its therapeutic potential in AP remains unclear. Here, we demonstrate that PHGDH overexpression or exogenous serine supplementation mitigates AP-associated elevations in serum digestive enzymes and reduces pancreatic histopathological injury. Notably, serine administration exerts pronounced therapeutic benefits in SAP. Previous studies reported that a chow diet containing 10% serine alleviates post-diabetic pancreatitis, suggesting that the efficacy of serine supplementation in AP is highly dependent on administration route, dosage, and may vary across AP subtypes and severity grades ^39^. Beyond animal models, we successfully established pancreatic organoids from human iPSC using an optimized protocol ^45^, enabling evaluation of serine’s protective effects in a human-relevant system. Collectively, these findings provide a strong preclinical foundation for the clinical translation of serine-based therapies in AP.

### Serine restored metabolic dysregulation and alleviated oxidative stress in acute pancreatitis

Oxidative stress plays a central role in multiple forms of pancreatic acinar cell death in AP, including necrosis ^46,47^, apoptosis ^46,48,49^, necroptosis ^49^ and ferroptosis ^50–52^. Serine has been reported to exert antioxidant and anti-inflammatory effects ^16,39–44^. Consistent with this, our subsequent single-cell analyses revealed that serine increases the proportion of normal acinar cells and reverses the expression of oxidative stress-related genes. Further *in vitro* assays and three-dimensional human pancreatic organoid models demonstrated that serine not only markedly attenuates acinar cell necrosis, but also reduces ROS levels and promotes the synthesis of cysteine and GSH. Both cysteine and GSH are critical for counteracting oxidative stress and inhibiting inflammatory cascades in AP ^53–60^. We next investigated whether serine directly fuels GSH synthesis through *de novo* cysteine production. Surprisingly, tracing experiments using either [^13^C_3_-serine] or [^13^C_6_-glucose] showed that less than 5% of cysteine is generated directly from this process, suggesting that serine contributes minimally to *de novo* cysteine biosynthesis and may regulate cysteine homeostasis via indirect or non-canonical mechanisms. Recent studies indicate that cysteine in pancreatic tissue arises from both *de novo* biosynthesis and extracellular uptake ^20,61,62^. Accordingly, [^13^C_6_-cystine] labeling revealed rapid intracellular cysteine generation in acinar cells, and this conversion was significantly accelerated by serine in our study. These findings indicate that cystine transport is impaired following acinar cell injury, whereas serine enhances this process.

Cysteine depletion is a hallmark of AP and a major contributor to GSH loss and oxidative stress ^63^. Although supplementation with cysteine precursors such as N-acetylcysteine (NAC) has shown protective effects in preclinical models by restoring GSH levels and reducing ROS, clinical trials have yielded inconsistent or disappointing results ^54,64,65^. One potential explanation is that NAC efficacy depends on cellular uptake mechanisms. Extracellular cysteine is unstable and rapidly oxidized to cysteine, whose uptake depends on the SLC7A11 transporter ^66^. Thus, supplementation alone may be insufficient when SLC7A11 function is compromised. Our study might contribute to the clinical antioxidant synergistic therapy of AP.

SLC7A11, a key cystine influx transporter on the acinar cell membrane, is degraded in SAP models, exacerbating oxidative stress and accelerating ferroptosis ^26–30^. Our metabolic flux analyses provide a mechanistic explanation for the pronounced impact of SLC7A11 impairment on cellular oxidative stress and injury. Importantly, serine stabilizes SLC7A11 protein *in vitro* and *in vivo*, and its antioxidant effect is abolished in *Slc7a11* knockout mice, underscoring the essential role of SLC7A11 in mediating serine’s protective effects. Elucidating how serine regulates SLC7A11 stability is therefore critical to understanding its antioxidant mechanism in AP.

### The serine-NEDD4-SLC7A11 ubiquitination regulatory network

SLC7A11 is the functional light-chain subunit of system x-c, a sodium-independent, chloride-dependent anionic L-cystine/L-glutamate antiporter on the cell surface ^62^. Its expression and activity are regulated at multiple levels, including transcriptional control, epigenetic modification, protein stability, interactions with other proteins, post-transcriptional and post-translational regulation, and transporter activity ^67,68^. In the present study, both SLC7A11 mRNA and protein levels were significantly reduced in AP models *in vivo* and *in vitro*. Notably, serine supplementation did not alter SLC7A11 mRNA expression but markedly increased its protein abundance and attenuated TLCS-induced SLC7A11 ubiquitination in PACs and 266-6 cells. SLC7A11 protein stability is controlled by the ubiquitin-proteasome system, in which E3 ubiquitin ligases attach ubiquitin chains to target proteins for degradation, while deubiquitinases remove ubiquitin moieties to stabilize substrates. More than ten E3 ligases have been reported to target SLC7A11, including SOCS2 ^24,31,69^, SOCS6 ^70^, TRIM3 ^71^, WWP1 ^72^, FBXW7 ^73^, NEDD4L ^74^, TRIM21 ^75,76^, TRIM7 ^32^, TRIM26 ^77^, HECTD3 ^78,79^, HRD1 ^80^ and KCTD10 ^81^. Similarly, multiple deubiquitinases stabilize SLC7A11, such as USP14 ^82^, USP18 ^83^, OTUB1 ^84–88^,OTUD5 ^89^ stabilize SLC7A11. Therefore, SLC7A11 ubiquitination appears to be tissue-and disease-specific.

Through IP-MS and Co-IP in acinar cells, we identified NEDD4 as a previously unreported E3 ubiquitin ligase mediating SLC7A11 ubiquitination in AP, expanding the tissue-specific regulatory landscape of SLC7A11. LiP-MS analysis of acinar cell lysates revealed that serine directly binds NEDD4, rather than SLC7A11. NEDD4, a representative member of the HECT family of E3 ubiquitin ligases, mediates substrate-specific ubiquitination via recognition of PY motifs or adaptor proteins ^90,91^. It contains an N-terminal WW domain for substrate recognition and a C-terminal HECT domain that catalyzes ubiquitin transfer ^92,93^. The HECT domain consists of an N-lobe and a C-lobe connected by a flexible hinge loop, which enables the C-lobe to accept ubiquitin from an E2 enzyme and transfer it to the substrate. LiP-MS of recombinant human NEDD4 demonstrated that serine binds directly to the HECT hinge loop, inducing conformational changes that modulate its catalytic activity. Complementary interaction studies confirmed robust binding between serine and NEDD4, with dissociation equilibrium constants (KD) of 0.46 μM by MST and 9.278 μM by SPR. In summary, our results identify NEDD4 as a novel E3 ligase for SLC7A11 in AP and reveal the HECT-domain hinge loop as a critical regulatory site. Serine directly binds NEDD4 with high affinity, modulating its enzymatic activity and thereby stabilizing SLC7A11.

### Expanding functional roles beyond acute pancreatitis: the serine-NEDD4-SLC7A11 axis in the pancreatic pathologies and beyond

Previous studies have linked serine metabolism to SLC7A11 regulation in several disease contexts, although these observations have largely been interpreted through the lens of metabolic enzyme activity within the *de novo* serine synthesis pathway. In diabetic kidney disease, PHGDH enhances SLC7A11 expression by stabilizing Y-box binding protein 1 in podocytes ^94^. In bladder cancer, PHGDH promotes SLC7A11 expression through stabilization of poly(rC)-binding protein 2 ^95^, while in colorectal cancer, phosphoserine aminotransferase 1 deficiency reduces SLC7A11 expression ^96^. Notably, the relationship is bidirectional: SLC7A11 can influence serine metabolism. Under cystine-limited conditions, impaired SLC7A11 function promotes glucose uptake and activates PHGDH-dependent serine biosynthesis via the transsulfuration pathway to preserve cysteine production and redox balance ^33^. In metabolic-associated steatotic liver disease, excessive hepatic SLC7A11 activity drives serine depletion through enhanced GSH synthesis, thereby impairing *de novo* cysteine production and exacerbating ferroptosis, which can be reversed by serine supplementation ^97^. Together, these findings define the current understanding of serine–SLC7A11 crosstalk as an indirect metabolic circuit governed primarily by enzymatic regulation.

Beyond this established metabolic interplay, a broader and mechanistically unresolved pattern has emerged. Across sterile inflammatory diseases in multiple organs, serine deficiency and SLC7A11 downregulation consistently co-occur, suggesting a functional relationship that extends beyond conventional metabolic interdependence. In drug-induced and toxic liver injury, reduced serine levels are accompanied by suppression of SLC7A11, contributing to ferroptotic cell death ^98,99^, while in metabolic liver disease, the glutamate-to-(serine+glycine) ratio correlates with fibrosis severity ^100^. Similar patterns are observed in ischemia–reperfusion injury across several organs. In ischemia–reperfusion injury, this convergent pattern extends across multiple organs. In cerebral ischemia–reperfusion, PHGDH downregulation in astrocytes drives oxidative stress and pyroptosis ^101^, while SLC7A11 is simultaneously suppressed ^102–106^. In hepatic ischemia–reperfusion, SHMT2 deficiency aggravates tissue injury and is associated with reduced SLC7A11 expression ^107 108^. Likewise, SLC7A11 is downregulated in myocardial ischemia–reperfusion ^105,109–111^, whereas in renal and intestinal ischemia–reperfusion, modulation of SLC7A11 critically determines ferroptotic sensitivity and tissue injury ^112–114^. Yet, despite this extensive correlative evidence, no prior study has addressed whether serine can directly modulate SLC7A11 protein abundance or function—a critical gap that has left the causal directionality of this relationship unresolved.

Here, we bridge this gap by uncovering a previously unrecognized signaling mechanism: serine directly binds the E3 ubiquitin ligase NEDD4, interfering with NEDD4-mediated ubiquitination and degradation of SLC7A11. This interaction stabilizes SLC7A11 protein and promotes cystine transport in pancreatic acinar cells. NEDD4 acts as a central regulator in this network. The relevance of NEDD4 in pancreatic biology extends well beyond acinar homeostasis. In β cells, it interacts with the ubiquitin-conjugating enzyme E2 E2 to promote polyubiquitin chain formation and inhibit proliferation ^115^. In pancreatic cancer, NEDD4 drives tumor progression by degrading phosphatase and tensin homolog and activating the phosphoinositide 3-kinases / protein kinase B pathway ^116^, suggesting that its inhibition may represent a therapeutic strategy across pancreatic diseases.

The therapeutic implications of the serine–NEDD4–SLC7A11 axis must, however, be interpreted in light of the pronounced cell-type specificity and functional pleiotropy of its downstream components. In pancreatic acinar cells, SLC7A11 upregulation enhances cystine uptake, elevates cysteine, fortifies antioxidant defenses, and prevents stress-induced injury and ADM ^20^. In β cells and islet-resident macrophages, SLC7A11 supports GSH homeostasis ^117,118^. By contrast, in pancreatic ductal adenocarcinoma, elevated SLC7A11 suppresses ferroptosis, promotes metabolic adaptation, and drives tumor progression, correlating with poor prognosis and chemoresistance ^119–122^. Serine itself exhibits similarly context-dependent effects: while elevated serine levels and PHGDH upregulation fuel tumor growth and metastasis ^123–125^, serine supplementation exerts unequivocal protection in inflammatory and oxidative injury settings ^16,39–44^. This dual nature underscores the need for cell-type-selective strategies that can harness the protective effects of the serine–NEDD4–SLC7A11 axis in inflammatory pathologies without inadvertently promoting neoplastic progression. Collectively, these findings establish the serine-NEDD4-SLC7A11 axis as a critical regulator of acinar cell homeostasis in AP and suggest its broader relevance in other pancreatic disorders, tumors and sterile inflammatory diseases.

### Limitations of the study

Several limitations of this study should be recognized. Firstly, whether the mechanism by which serine stabilizes SLC7A11 operates in other sterile inflammatory diseases, such as drug-induced and toxic liver injury, ischemia–reperfusion injury and diabetes remains to be established. Secondly, defining the role of NEDD4 in regulating the ubiquitination of other proteins will help assess the therapeutic specificity of NEDD4 inhibition in AP. Thirdly, the direct regulatory role of serine on NEDD4 remains to be further confirmed via site-directed mutagenesis or knockout in mice. Fourthly, the function and regulatory mechanisms of the *Ccr2* high inflammatory acinar cell subset—a novel population identified by scRNA-seq—require further elucidation. Finally, the clinical translational value of serine for diagnosis and treatment still requires validation through multicenter studies.

## Methods

### Materials and Reagents

Resources, identifiers, and other relevant details for all materials and reagents are listed in **Supplementary Tab. 9**. **AP patients’ sample collection and data reanalysis** Human AP patients’ sample data used in this study were obtained from a prior study ^35^, with relevant patient information and ethical approval documented in the original report ^35^. Further reanalysis of these datasets was performed in the present study.

### Animal studies

Male C57BL/6 mice (6–8 weeks old, 20–25 g) were purchased from Jiangsu Jicui Yaokang Biotechnology Co., Ltd. (Jiangsu, China). All animals were housed in the specific pathogen-free (SPF) barrier facility at the Experimental Animal Center of West China Hospital, Sichuan University, under controlled conditions (temperature 22 ± 2°C, relative humidity 50–60%, 12-h light/dark cycle). Mice had free access to standard rodent chow and sterilized drinking water, with 5 mice per cage. All animal procedures were approved by the Institutional Animal Care and Use Committee of West China Hospital, Sichuan University (Approval numbers: 2021042A, 20260123004).

To achieve specific *Phgdh* gene modulation in pancreas, recombinant adeno-associated virus (AAV) was administered via retrograde pancreatic duct injection. For *Phgdh* knockdown: Mice received 100 μL of AAV carrying short hairpin RNA targeting *Phgdh* (AAV-sh*Phgdh*) or negative control virus (AAV-NC) via retrograde pancreatic duct injection. For *Phgdh* overexpression: Mice received 100 μL of AAV carrying the *Phgdh* gene (AAV-*Phgdh*; titer 5 × 10¹² VG/mL) or AAV-NC via retrograde pancreatic duct injection. Following viral injection, mice were allowed to recover for 4 weeks to ensure stable transgene expression in the pancreas before induction of AP.

Male global *Slc7a11* knockout (*Slc7a11* KO) mice were purchased from Jiangsu Jicui Yaokang Biotechnology Co., Ltd. (Jiangsu, China). All animals were housed in the SPF barrier facility at the Experimental Animal Center of West China Hospital, Sichuan University, under controlled conditions (temperature 22 ± 2°C, relative humidity 50–60%, 12-h light/dark cycle). Mice had free access to standard rodent chow and sterilized drinking water, with 5 mice per cage. All animal procedures were approved by the Institutional Animal Care and Use Committee of West China Hospital, Sichuan University (Approval numbers: 20260123003).

### Cell lines

PACs were isolated from male C57BL/6J mice between the age of 6–8 weeks old by collagenase digestion as previously reported ^126^.

The mouse pancreatic acinar carcinoma cell line 266-6 was obtained from the American Type Culture Collection (ATCC). Cells were cultured in DMEM supplemented with 10% fetal bovine serum (FBS) and 1% penicillin-streptomycin at 37°C in a 5% CO_2_ humidified incubator. Lentiviral particles carrying Flag-*Slc7a11* or negative control (WT) were purchased from Shanghai Genechem Co., Ltd. (Shanghai, China). Cells were seeded at an appropriate density 24 h prior to infection. After 16 h, medium was replaced with complete medium. At 36 h post-infection, EGFP expression was assessed by fluorescence microscopy, and stable Flag-SLC7A11-overexpressing clones were selected using puromycin. Plasmids carrying *Nedd4* (Shanghai Genechem Co., Ltd.) or empty vector control were transfected into 266-6 cells using Lipofectamine 3000. Cells were seeded 24 h prior to transfection and harvested 48 h post-transfection for downstream analyses.

### Pancreatic organoid culture

Human induced pluripotent stem cell (iPSC)-derived pancreatic organoids were generated using a Matrigel-free protocol to induce branching morphogenesis and acinar-trunk patterning ^45^. Human iPSC (human iPSC-F1, female; CellAPY Biotechnology Co., Ltd., Beijing, China) at passage 25 were cultured on 1% Matrigel-coated (Corning, New York, USA) germfree 6-well plates (Thermo Fisher Scientific, Waltham, USA) at 37°C, 5% CO_2_. Cultures were maintained and replaced daily with mTeSR1 medium (STEMCELL Technologies, Vancouver, Canada) with 1% penicillin-streptomycin. Human iPSC at approximately 70% confluency were dissociated into single cells using Gentle Cell Dissociation Reagent (STEMCELL Technologies, Vancouver, Canada) supplemented with 20 μM ROCK inhibitor (Y-27632, MCE, Shanghia, China). Detailed procedures were described by ^45^. Briefly, human iPSC were differentiated into pancreatic progenitors under two-dimensional culturing for 10 days using Pancreatic Organoid Culture Kit (NVM11011, Beijing Novital Biotechnology Ltd., Beijing, China). At day 10, progenitors were dissociated with TrypLE (Gibco, New York, USA), resuspended and seeded at 1,5000-2,0000 cells/microwell in aggrewell plates (24 wells, STEMCELL Technologies, Vancouver, Canada). At day 12, organoids were transferred into 6 well plate of ultra-low attachment surface and placed on an orbital shaker. Medium from culture kit was changed daily until day 15. Organoids were collected at D16 for experiments and analysis.

### AP models and interventions

To induce mild edematous AP model, male C57BL/6J mice (6–8 weeks old, ∼22 g) were administered 7 intraperitoneal injections of cerulein (CER, 50 μg/kg) at hourly intervals. The time points of injection were set at 0, 1, 2, 3, 4, 5 and 6 h. For serine pharmacological intervention, serine was administered intraperitoneally at 0, 4, and 8 h (1.6 g/kg per dose). And serum and pancreatic tissue were collected at 12 h. For heclin pharmacological intervention, heclin was administered intraperitoneally at 3 and 6 h (1 mg/kg and 5 mg/kg per dose, respectively), and serum and pancreatic tissue were collected at 12 h.

To induce severe necrotizing AP model, mice (8 weeks old, ∼25 g) received 2 intraperitoneal injections of L-arginine (ARG, 4 g/kg, pH 7.0) at hourly intervals. The time points of injection were set at 0 and 1 h. For serine pharmacological intervention, serine was administered intraperitoneally daily at 4, 8, 24,28,32, 48, 52, and 56h (1.6 g/kg per dose) and serum and pancreatic tissue were collected at 72 h.

The sample collection process is as follows: At predetermined time points, mice were euthanized, and blood and tissues were collected. Blood was obtained via retro-orbital bleeding, allowed to clot at room temperature for 30 min, centrifuged at 1500 g for 10 min, and the supernatant re-centrifuged at 1500 g for 5 min. The resulting serum was aliquoted and stored at-80°C for biochemical and oxidative stress assays. The pancreas was divided into head, body, and tail portions: the head for MPO activity; the body near the tail fixed in neutral formalin for hematoxylin-eosin (H&E) staining and immunohistochemistry; the mid-body snap-frozen in liquid nitrogen (divided into 3 aliquots) for metabolomics, trypsin activity, or western blot analysis; and the tail minced in RNA later for real-time quantitative polymerase chain reaction (RT-qPCR). All samples were stored at-80°C.

### Detection of AP severity indicators and oxidative stress indicators

Serum amylase and lipase activity, pancreatic MPO activity, and pancreatic trypsin activity as previously described ^36,126^. Paraffin-embedded pancreatic sections were subjected to hematoxylin and eosin (H&E) staining, immunohistochemistry and immunofluorescence. Established scoring criteria were used for pancreatic histopathology assessment, including pancreatic edema, inflammatory cell infiltration, and cell necrosis ^127^.

Superoxide dismutase (SOD) and glutathione peroxidase (GPx) activities in PACs, as well as ATP concentrations in PACs and pancreatic organoids, were quantified using commercial assay kits from Nanjing Jiancheng Bioengineering Institute according to the manufacturer’s instructions.

ROS levels in paraffin-embedded mouse pancreatic sections were detected using a commercial assay kit (BB-470513), as detailed in our previous study ^128^

Detailed methods for western blotting (WB) and real-time quantitative PCR (RT-qPCR) are provided in our previous studies ^36^. Primer sequences for the target genes are listed in Table

### Transmission electron microscopy (TEM)

Fresh tissue samples were carefully dissected to minimize mechanical damage, and tissue blocks were cut to dimensions not exceeding 1 mm × 1 mm × 1 mm. Samples were immediately fixed in electron microscopy fixative at 4°C for 2-4 h, followed by three washes with 0.1 M phosphate buffer (PB, pH 7.4) for 15 min each. Samples were post-fixed with 1% osmium tetroxide in 0.1 M PB (pH 7.4) at room temperature for 2 h, then washed three times with 0.1 M PB (pH 7.4) for 15 min each. Dehydration was performed through a graded ethanol series (50%, 70%, 80%, 90%, 95%, and 100%) followed by 100% acetone and 100% acetone, with 15 min per step. Samples were infiltrated with acetone: Spurr’s resin mixtures (1:1 for 2–4 h, then 2:1 overnight), followed by pure Spurr’s resin for 5–8 h. Samples were then embedded in pure Spurr’s resin and polymerized at 60°C for 48 h. Ultrathin sections (60–80 nm) were cut using an ultramicrotome and stained with uranyl acetate and lead citrate (15 min each). Sections were air-dried overnight at room temperature and examined under a transmission electron microscope. Images were captured for subsequent analysis.

### Targeted metabolomics, individual concentration perturbation profile (ICPP) calculation and disease-specific pathways identification

The targeted metabolomics data of plasma samples from clinical patients were derived from our previously published study ^35^. Targeted metabolomics analysis was also performed on serum samples from CER-injured mice at different time points (0, 1, 4, 8, 12, and 24 h), pancreatic tissues from CER-injured mice treated with serine (1.6 g/kg i.p. at 0, 4, and 8 h; sampled at 12 h), and ARG-injured mice at different time points (0, 8, 24, 48, and 72 h). Detailed protocols for sample collection, preparation, chromatographic separation, mass spectrometric detection, and metabolite quantification are described in our previous study ^36^.

Raw metabolomics data were preprocessed as follows: metabolites with missing values in >10% of samples were excluded; remaining missing values were imputed using the median concentration of the respective metabolite across all samples. For quality control, metabolites exhibiting a relative standard deviation (RSD) >15% in pooled quality control samples were removed. Pancreatic tissue metabolite concentrations were normalized to total protein content, as measured using a bicinchoninic acid protein assay kit. All concentration values were then log₂-transformed to approximate normality and stabilize variance prior to downstream analysis. The significant metabolites were analyzed as described in our previous study ^36^. After data preprocessing, univariate and multivariate statistical analyses were performed, including PCA and orthogonal partial least squares-discriminant analysis (OPLS-DA), to generate parameters including *P*-value, FC, and variable importance in projection (VIP) score. Metabolites with FC > 1.5 or FC < 0.67, *P*-value < 0.05, and VIP > 1 were defined as differential metabolites.

We conducted a metabolic pathway perturbation analysis on the serum of CER-AP model, and the pancreas of ARG-AP model, and previously generated human plasma targeted metabolomics dataset ^129^.

For each metabolite *i* in an individual subject *j*, we calculated a Z-score to quantify its deviation from the reference distribution in the control group. The Z-score was computed as:

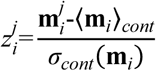

Where **m_i_***^j^* is the concentration of metabolite *i* in subject *j*, 〈**m***_i_*〉*_cont_* is the mean concentration in control subjects, and *σ_cont_*(**m***_i_*) is the standard deviation. If a subject belonged to the control group, it was excluded from the reference distribution calculation. Metabolites with |*z^j^j* |≥z_thresh_ (where z_thresh_ = 2.5) were considered significantly perturbed in that individual. An ICPP was thereby generated for each subject, serving as a metabolic “barcode” indicating up-regulated (*z^j^*>z_thresh_) or down-regulated (*z^j^*<-z_thresh_) metabolites relative to controls.

For each subject’s ICPP, pathway interruption was performed using Fisher’s exact test (hypergeometric distribution) to assess over-representation of perturbed metabolites within KEGG pathways. *P*-values were adjusted for multiple comparisons using the Holm-Bonferroni method. Pathways were deemed significantly enriched if the adjusted *P*-value < 0.01. To identify pathways with consistent relevance to AP, the frequency of each pathway was quantified across all case subjects. Pathways exhibiting high frequency (indicating homogeneity among cases) were selected as potential disease-specific pathways for further validation.

### Absolute quantitation of serine in serum/plasma and pancreatic tissue

A mixture consisting of 20 μL serum/plasma and 100 μL methanol was vortexed for 5 min, incubated at −80 °C for 30 min, sonicated for 10 min, and then centrifuged at 16,000 ×g and 4 °C for 20 min. The supernatant was filtered through a 0.22 μm membrane. A standard calibration curve for serine was constructed by spiking gradient concentrations into 5% BSA blank matrix. For pancreatic tissue samples, approximately 4 mg of tissue was homogenized in 500 μL of 80% (v/v) methanol. Following vortexing, sonication, and centrifugation, the supernatant was collected, dried, reconstituted in 60% (v/v) acetonitrile, and filtered through a 0.22 μm membrane. The standard curve of serine in tissues was established using an isotope substitution method, in which gradient-diluted [^13^C_3_]-serine was spiked into control pancreatic tissue samples. The standard curve of serine in tissues was established using an isotope substitution method, in which gradient-diluted [^13^C_3_]-serine was spiked into control pancreatic tissue samples. The intensity of [^13^C_3_]-serine was used to calculate the serine concentration. LC-MS/MS analysis was carried out on a Nexera LC-30A UHPLC system coupled with an AB Sciex QTRAP 5500+ mass spectrometer (AB Sciex, USA). Chromatographic separation was performed on an Acquity BEH Amide Column (2.1 × 100 mm, 1.7 μm; Waters, USA). Mobile phase A consisted of 5% (v/v) acetonitrile supplemented with 5 mM ammonium acetate and 0.1% (v/v) acetic acid, while mobile phase B comprised 95% (v/v) acetonitrile with the same additives. Column temperature and flow rate were set at 40 °C and 0.4 mL/min, respectively. The gradient program was set as follows: 94% B at 0 min, 94% B at 1 min, 50% B at 5 min, 50% B at 6 min, 94% B at 7 min, and 94% B at 17 min. In positive ionization mode with multiple reaction monitoring (MRM), serine was detected using the transition of *m/z* 106.0 → 60.1, with a declustering potential (DP) of 50 eV and a collision energy (CE) of 15 eV. [^13^C_3_]-serine was detected using the transition of *m/z* 109.0 → 62.1 (DP 50 eV, CE 15 eV). The injection volume was 5 μL and data were extracted and quantified using SCIEX OS software.

### Quantitation of cysteine and GSH in pancreatic acinar cells

GSH and cysteine were determined using an NEM derivatization method. PACs were washed with HEPES buffer, and then 1 mL of 80% methanol containing 12.5 mM NEM was added. The mixture was vortexed, kept at-80°C for 30 min, sonicated, and centrifuged. The supernatant was dried, reconstituted in 50% (v/v) methanol, and filtered through a 0.22 μm membrane. 5 μL of the supernatant was injected into the Nexera LC-30A UHPLC system coupled to an AB Sciex QTRAP 6500+ mass spectrometer (AB Sciex, USA). An Acquity BEH Amide column (2.1 × 100 mm, 1.7 μm; Waters, USA) was used for separation. Mobile phase A and B consisted of 10% and 90% acetonitrile (v/v), respectively, both supplemented with 10 mM ammonium acetate and 0.2% acetic acid. The column oven was kept at 40 °C with a flow rate of 0.3 mL/min. The gradient elution program was set as: 90% B from 0 to 1.5 min, decreased to 45% B at 5 min and held until 10 min, then returned to 90% B at 12 min and maintained until 25 min. Quantification was performed in positive MRM mode with the ion transitions of *m/z* 247.0 → 230.1 (DP 50 eV, CE 17 eV) for cysteine derivative and *m/z* 433.4 → 304.1 (DP 85 eV, CE 21 eV) for GSH derivative. The data processing and quantification were carried out using SCIEX OS software.

### Mass spectrometry imaging (MSI) analysis

MSI analysis of pancreatic tissues followed the procedures described in our previous work ^35^. Mass spectrometry data were collected with Xcalibur software (Version 2.2, Thermo Scientific, USA). Raw data files were converted to the.cdf format via the Xcalibur file conversion tool, and the resulting.cdf files were imported into Massimager Pro (Version 1.0, China) for the reconstruction of tissue ion images. After subtraction of background ions, three regions of interest (ROIs) were randomly selected from each tissue section using a standard 5 × 5 pixel rectangular frame. Each ROI yielded an independent mass spectral profile, which contained m/z and intensity information and was saved as a.txt file. Finally, the signal intensities of serine (m/z 104.0351) were quantified, respectively.

### Single-cell RNA sequencing (scRNA-seq) and bioinformatic analysis

Pancreas were transported in sterile culture dish with 10 ml 1x Dulbecco’s Phosphate-Buffered Saline (DPBS) on ice to remove the residual tissue storage solution, then minced on ice. We used dissociation enzyme 0.25% Trypsin and 10 μg/mL DNase I dissolved in PBS with 5% FBS to digest the tissues. Pancreas were dissociated at 37℃ with a shaking speed of 50 r.p.m for about 40 min. We repeatedly collected the dissociated cells at interval of 20 min to increase cell yield and viability. Cell suspensions were filtered using a 40 μm nylon cell strainer and red blood cells were removed by 1X Red Blood Cell Lysis Solution. Dissociated cells were washed with 1x DPBS containing 0.4% FBS. Cells were stained with 0.4% AO/PI to check the viability on Countstar Rigel S2 (Countstar, Shanghai). The sample was then sent to Majorbio Bio-pharm Technology Co.,Ltd (Shanghai, China) for testing.

Beads with unique molecular identifier (UMI) and cell barcodes were loaded close to saturation, so that each cell was paired with a bead in a Gel Beads-in-emulsion (GEM). After exposure to cell lysis buffer, polyadenylated RNA molecules hybridized to the beads. Beads were retrieved into a single tube for reverse transcription. On cDNA synthesis, each cDNA molecule was tagged on the 3’end (that is, the 5’end of a messenger RNA transcript) with UMI and cell label indicating its cell of origin. Briefly, 10x beads that were then subject to second-strand cDNA synthesis, adaptor ligation, and universal amplification. Sequencing libraries were prepared using randomly interrupted whole transcriptome amplification products to enrich the 3’ end of the transcripts linked with the cell barcode and UMI. All the remaining procedures including the library construction were performed according to the standard manufacturer’s protocol (Chromium Single Cell 3ʹ v3.1). Sequencing libraries were quantified using a High Sensitivity DNA Chip (Agilent) on a Bioanalyzer 2100 and the Qubit High Sensitivity DNA Assay (Thermo Fisher Scientific, USA). The libraries were sequenced on NovaSeq X Plus /DNBSEQ platform using 2×150 chemistry. The sequencing and bioinformatic analysis were performed on platform of Majorbio Co., Ltd (Shanghai, China).

Reads were processed using the Cell Ranger V7.1.0 pipeline with default and recommended parameters. FASTQs generated from Illumina sequencing output were aligned to the Novaseq genome, version GM4563, using the STAR algorithm ^130^. Next, Gene-Barcode matrices were generated for each individual sample by counting UMIs and filtering non-cell associated barcodes. Finally, we generate a gene-barcode matrix containing the barcoded cells and gene expression counts. This output was then imported into the Seurat (v4.1.1) R toolkit for quality control and downstream analysis of our scRNA-seq data ^131^. All functions were run with default parameters, unless specified otherwise. We first filtered the matrices to exclude low-quality cells using a standard panel of three quality criteria:

(1) number of detected transcripts (number of unique molecular identifiers); (2) detected genes; and (3) percent of reads mapping to mitochondrial genes (The numbers out of the limit of mean value +/-2-fold of standard deviations). The expression of mitochondria genes was calculated using PercentageFeatureSet function of the Seurat package [3]. The normalized data (NormalizeData function in Seurat package) was performed for extracting a subset of variable genes. Variable genes were identified while controlling for the strong relationship between variability and average expression. Next, we integrated data from different samples after identifying ‘anchors’ between datasets using FindIntegrationAnchors and IntegrateData in the Seurat package ^131,132^. Then we performed PCA and reduced the data to the top 30 PCA components after scaling the data. We visualized the clusters on a 2D map produced with t-distributed stochastic neighbor embedding (t-SNE).

Cells were clustered using graph-based clustering of the PCA reduced data with the Louvain Method after computing a shared nearest neighbor graph [3]. For sub-clustering, we applied the same procedure of scaled, dimensionality reduction, and clustering to the specific set of data (usually restricted to one type of cell). For each cluster, we used the Wilcoxon Rank-Sum Test to find significant deferentially expressed genes comparing the remaining clusters. SCINA ^133^ and known marker genes were used to identify cell type.

To identify DEGs (differential expression genes) between two different samples or clusters, was performed using the function FindMarkers in Seurat,using a Wilcoxon rank-sum test. Essentially, DEGs with |log_2_FC|>0.25 and q value ≤ 0.05 were considered to be significantly different expressed genes. In addition, functional-enrichment analysis of GO and KEGG were performed to identify which DEGs were significantly enriched in GO terms and metabolic pathways at Bonferroni-corrected *P*-value ≤ 0.05 compared with the whole transcriptome background. GO functional enrichment analysis were carried out by Goatools (https://github.com/tanghaibao/Goatools). KEGG functional enrichment analysis were carried out by Python scipy software.

### Proteomics and bioinformatic analysis

For serine-treated CER model mice, pancreatic samples were harvested and subjected to proteomic analysis. Detailed protocols for sample preparation, chromatographic separation, mass spectrometric detection, protein quantification, and data analysis have been described in our previous study ^134^. Differentially expressed proteins were screened with a *P*-value < 0.05 and FC > 1.5 or FC < 0.67. These proteins were further annotated using GO and KEGG enrichment analysis.

### Isolation of pancreatic acini and *in vitro* studies

PACs were freshly prepared by collagenase digestion. Acinar cell necrosis, ROS accumulation, and GSH levels were assessed using PI/hoechst staining, H₂DCFDA fluorescence, and a monochlorobimane probe, respectively, as previously described ^11,128,135^. PACs were stimulated to induce injury using TLCS (500 μmol/L), ARG (20 mmol/L) and H_2_O_2_ (100 μmol/L) as model groups. PACs were incubated with serine (100 mmol/L), heclin (0.1, 1, 2 and 5 μmol/L) for 30 min, followed by co-incubation with the modeling agents for 30 min.

### Cell metabolic flux analysis

PACs for the [^13^C_6_]-glucose tracing, [^13^C_3_]-serine tracing and [^13^C_6_]-cystine tracing were prepared in the same way as previously described ^128^, but the HEPES buffer contained certain stable isotope-labeled metabolites ([^13^C_6_]-glucose, 10 mM; [^13^C_3_]-serine, 400 μM; [^13^C_6_]-cystine, 100 μM). [^13^C_6_]-cystine was generated from [^13^C_3_]-cysteine by oxidation with H_2_O_2_, according to a previously reported method ^61^. PACs were isolated freshly and divided into three groups (Ctrl, AP, and APS) with 3 or 4 replicates. Before extraction, cells were fed with [^13^C_6_]-glucose for 2 h, or [^13^C_3_]-serine for 30 min, or [^13^C_6_]-cystine for 5 min. Metabolites were extracted as previously described ^11^. LC-MS analysis was performed using a Vanquish UHPLC system coupled with a Q-Exactive Plus Orbitrap high-resolution mass spectrometer (Thermo Scientific, USA). The chromatographic column was an Acquity BEH Amide Column (2.1 × 100 mm, 1.7 μm, Waters, USA). Mobile phases A and B consisted of 5% and 95% acetonitrile (v/v), respectively, both containing 5 mM ammonium acetate and 0.1% (v/v) acetic acid. The column was maintained at 35°C with a flow rate of 0.3 mL/min. The gradient was programmed as: 94% B (0–1 min), 78% B at 7.5 min, 39% B (12–17 min), and 94% B (19–30 min). Data were acquired in positive ESI modes with full MS scans at 70,000 resolution. The injection volume was 5 μL. Instrumental control and data acquisition were achieved by Xcalibur software (version 4.2, Thermo Fisher Scientific, USA). Isotope correction for the extracted metabolite intensities was performed using the AccuCor1 R package with normal resolution correction.

### Ubiquitination Assay

To assess SLC7A11 ubiquitination, PACs and 266-6 cells were processed as follows: (1) PACs: Pre-treated with 100 mM serine for 30 min, stimulated with 500 μM TLCS for 30 min, and treated with MG132 (10 μM) for 4 h before harvest. (2) 266-6 cells: Flag-SLC7A11-overexpressing stable cells were treated with 100 mM serine and 500 μM TLCS for 24 h, with MG132 (10 μM) added for the final 4 h. Cells were lysed, and lysates were immunoprecipitated with anti-flag, anti-SLC7A11 or control IgG. Immunoprecipitates were washed, eluted, and analyzed by western blotting using anti-HA (for exogenous HA-tagged ubiquitin) or anti-ubiquitin antibodies. Input Vinculin served as controls.

### Immunoprecipitation-mass spectrometry (IP-MS)

To identify SLC7A11-interacting proteins, PACs were stimulated with 500 μM TLCS for 30 min and lysed. Lysate supernatant was divided: one portion served as input control, and the remainder was incubated overnight at 4°C with anti-SLC7A11 antibody or control IgG pre-coupled to agarose beads. After extensive washing, bound proteins were eluted, desalted, trypsin-digested, and analyzed on an Orbitrap Exploris 480 mass spectrometer (Thermo Scientific, USA). Data were processed using bioinformatics tools to identify SLC7A11-specific interactors.

### Co-immunoprecipitation (Co-IP)

The Co-IP assay was performed according to the kit instructions (KTD105, Abbkine). Briefly, lysates of PACs were immunoprecipitated in IP buffer containing IP antibody-coupled agarose beads, and protein-protein complexes were later analyzed by Western blot. Antibody for SLC7A11 (ab307601, Abcam) and NEDD4 (HA721191, HUABIO) was used. The labeled protein bands were visualized using a Chemiluminescent Detection System (Bio-Rad).

### Limited proteolysis-mass spectrometry (LiP-MS) analysis

TLCS-injured PACs treated with or without serine were washed with HEPES buffer and then lysed with lysis buffer. In contrast, the recombinant protein samples with or without serine supplementation were directly subjected to the reaction, bypassing the lysis step. The mixture was subjected to limited proteolysis with proteinase K (1:100 w/w, 25°C, 5 min). The reaction was terminated by heating at 98°C for 5 min. Subsequently, sodium deoxycholate (5% final), dithiothreitol (10 mM, 56°C, 1 h), and iodoacetamide (25 mM, 25°C, 30 min, dark) were sequentially added for deoxycholate addition, reduction, and alkylation, respectively. Following dilution with 100 mM ammonium bicarbonate to 1% deoxycholate, samples were digested first with lysyl-endopeptidase (1:100 w/w, 37°C, 2 h) and then with trypsin (1:100 w/w, 37°C, 16 h). Digestion was quenched with trifluoroacetic acid (10% v/v, pH < 3). Peptides were desalted on C_18_ microspin cartridges, eluted with 50% acetonitrile/0.1% trifluoroacetic acid, vacuum-dried, and reconstituted in 0.1% formic acid for subsequent LC-MS/MS analysis on an Orbitrap Exploris 480 mass spectrometer (Thermo Scientific, USA). Data-independent acquisition data were processed with Spectronaut 19.9. Prior to comparative analysis, data were median-normalized and filtered for proteotypic peptides.

### Microscale thermophoresis (MST) binding assay

The MST binding assay was performed as described previously ^136^. Recombinant NEDD4 protein was labeled using the Monolith Protein Labeling Kit RED-NHS (2nd Generation) and then mixed with serine at various concentrations. Serine was serially diluted twofold from an initial concentration of 100 μM. The mixture was loaded into capillaries and subjected to detection. Binding affinities between serine and NEDD4 were measured using a Monolith NT.115 instrument (NanoTemper, Germany). Dissociation equilibrium constants (KD) were calculated using the mass action equation in the NanoTemper software.

### Surface plasmon resonance (SPR) analysis

SPR experiments were performed on a Biacore X100 instrument (Cytiva). Experimental setups were designed using Biacore Control Software version 2.0.2, and all data were analyzed with Biacore Evaluation Software version 2.0.2.

For ligand immobilization, CM5 sensor chips were activated by injecting a 1:1 mixture of 0.4 M 1-ethyl-3-(3-dimethylaminopropyl)carbodiimide (EDC) and 0.1 M N-hydroxysuccinimide (NHS) (Cytiva, BR-1006-33) over flow cells 1 and 2 for 9 min. NEDD4 was immobilized on flow cell 2 by injecting 50 μg/ml recombinant NEDD4 diluted in 10 mM sodium acetate (pH 4.0; Cytiva, BR100352) until an immobilization level of approximately 15,310 RU was reached. Both flow cells were then blocked with 1 M ethanolamine (Cytiva, BR-1006-33) for 9 min. Flow cell 1 served as the reference surface.

Raw sensorgrams were imported into Biacore Evaluation Software 2.0.2. Data quality was assessed by examining baseline stability, binding levels, binding stability, and reference subtraction. Kinetic analysis was performed by fitting the sensorgrams to an appropriate binding model. Sensorgrams corresponding to the best-fit concentrations were exported for further analysis and visualization using third-party software (e.g., GraphPad Prism).

### Molecular docking

Based on LiP-MS results, molecular docking of serine to NEDD4 was conducted in Discovery Studio 2019. The initial structures of the NEDD4 HECT domain (PDB: 2XBF, http://www.rcsb.org/pdb) and serine (PubChem, https://pubchem.ncbi.nlm.nih.gov/) were downloaded and prepared by removing solvents and performing standard repairs (including residue correction and hydrogen addition). For flexible docking, the peptides identified by LiP screening were assigned as flexible residues, and the regulatory domain active site was set as the binding sphere. The highest-scoring docking pose was selected to represent the optimal binding mode.

## Statistical analysis

All data are presented as mean ± standard error of the mean (SEM). Statistical analysis was carried out by GraphPad Prism 10.1.2 software (La Jolla, CA, USA). Unpaired two-tailed Student’s t-test, one-way ANOVA with Dunnett’s multiple comparison tests, or non-parametric tests with the KruskaleWallis H test were used for data from two and several groups, respectively. *P*-value <0.05 was considered statistically significant.

### Reporting summary

Further information on research design is available in the Nature Portfolio Reporting Summary linked to this article.

## Data availability

scRNA-seq data have been uploaded to the Gene Expression Omnibus under accession numbers GSE327963.

## Code availability

No new algorithms were developed for this Article. Processed data and analysis code are available from the corresponding author.

## Acknowledgements

We acknowledged the funding from the National Natural Science Foundation of China (82170905 and 82522091 to D.D.); the Project of Sichuan Provincial Administration of Traditional Chinese Medicine Innovation Research Team (2023ZD04 to Q.X.); 1.3.5 project for disciplines of excellence, West China Hospital, Sichuan University to Q.X. We thank Rui Wang, Xiyu Wu, Qianlun Pu, Na Jiang, Wanmeng Li (Mass Spectrometry Center, Frontiers Science Center for Disease-related Molecular Network of West China Hospital, Sichuan University) for their assistance with MS analysis and technical support; Jingyao Zhang (Core Facilities of West China Hospital, Sichuan University) for her kind and professional help with the SPR assays in data acquisition and analysis; as well as Yue Li and Yi Zhang (Research Core Facility of West China Hospital, Sichuan University) for their support with histopathological examinations; Xiaoting Chen (Experimental Animal Center of West China Hospital, Sichuan University) for help with animal feeding; Yongjian Wen (West China Center of Excellence for Pancreatitis, Institute of Integrated Traditional Chinese and Western Medicine, West China Hospital, Sichuan University) for the design of *Phgdh* shRNA; Wei Huang and Juqin Yang (West China Center of Excellence for Pancreatitis and West China Biobank, West China Hospital, Sichuan University) for their assistance with human samples and quantification of metabolites. We also thank Chengdu Medical College for providing Discovery Studio software, and declare that the graphical abstract and some critical depicting images were created with BioR ender.com.

## Contributions

**Y. H.**: Investigation, Methodology, Formal analysis, Validation, Visualization, Data curation, Writing - original draft. **F. F.**: Investigation, Formal analysis, Validation, Visualization, Data curation, Writing - original draft. **L. D.**: Visualization, Data curation. **Y. W.**, **J. L.**, **J. Z.**, **J. Y.**, **Y. L.**, **M. W.** and **C. H.**: Methodology, Validation. **L. D.**: Resources. **P. L.**: Methodology. **H. C.**: Supervision. **J. D.**: Visualization, Data curation. **X. F.**: Supervision, Conceptualization. **Q. X.**: Supervision, Conceptualization, Funding acquisition, Resources. **D. D.**: Supervision, Conceptualization, Funding acquisition, Resources, Writing – review & editing

## Competing interests

The authors declare no competing interests.

## Declaration of generative AI and AI-assisted technologies in the writing process

During the preparation of this manuscript, ChatGPT (OpenAI) and DeepSeek were used to assist in language refinement. All outputs were carefully reviewed and revised by the authors, who assume full responsibility for the accuracy and integrity of the final manuscript.

## Supplementary figures and legends

**Supplementary Fig. 1:**
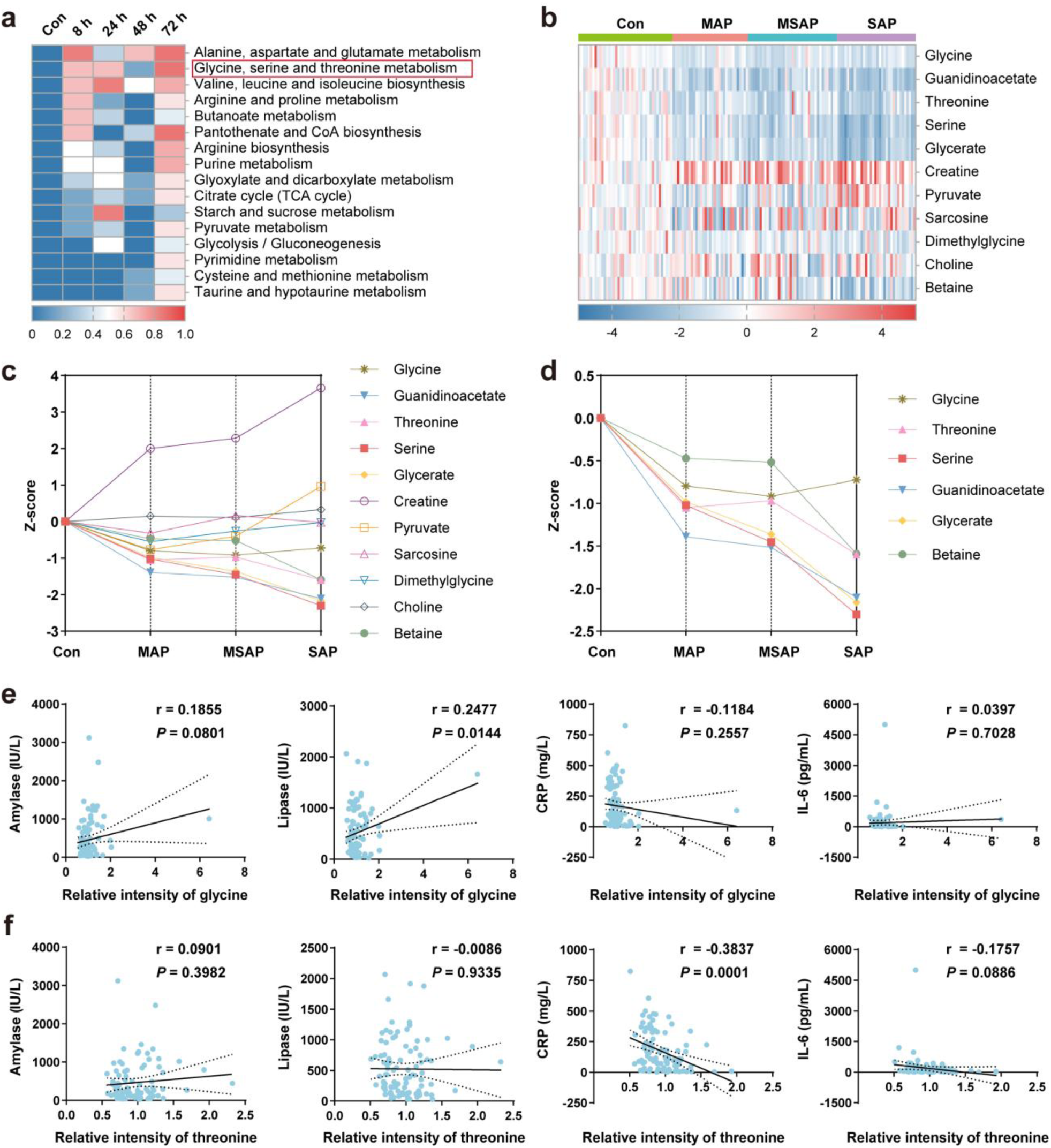
Disrupted glycine, serine and threonine metabolism in AP (related to. Fig. 1**). a,** Top 16 disease-specific metabolic pathways interrupted in serum of ARG-induced AP mice, identified by ICPP-based pathway perturbation analysis (adjusted *P* < 0.01). Targeted metabolomics were analyzed using pancreatic tissues from ARG-induced AP mice at five timepoints (8 h, 24 h, 48 h, 72 h after model establishment) and control mice (n=8). **b,** Heatmap showing the relative abundances of 11 metabolites in glycine, serine and threonine metabolism pathway across patients (day 1 after admission) with varying severities of AP and HCs. Rows represent individual metabolites; columns represent individual samples grouped by condition (AP vs HCs). Color scale indicates z-score values (red: upregulated in AP; blue: downregulated in AP; white: no change). **c,** Line graph illustrating the average z-score values of the 11 metabolites across AP patients and HCs, highlighting pathway-wide dysregulation in AP. Each line represents a metabolite; x-axis: groups; y-axis: normalized abundance or z-score. Data are presented as mean. **d,** Individual line graphs showing decreased levels of 6 out of the 11 metabolites in AP patients compared to HCs. Each line represents a metabolite; x-axis: groups (control vs. AP); y-axis: normalized abundance or z-score. Data are presented as mean. **e,** Pearson correlations of plasma glycine intensity with amylase, lipase, CRP and IL-6 levels (n = 90–97). **f,** Pearson correlations of plasma threonine intensity with amylase, lipase, CRP and IL-6 levels (n = 90–97). Values shown are mean ± SEM.

**Supplementary Fig. 2:**
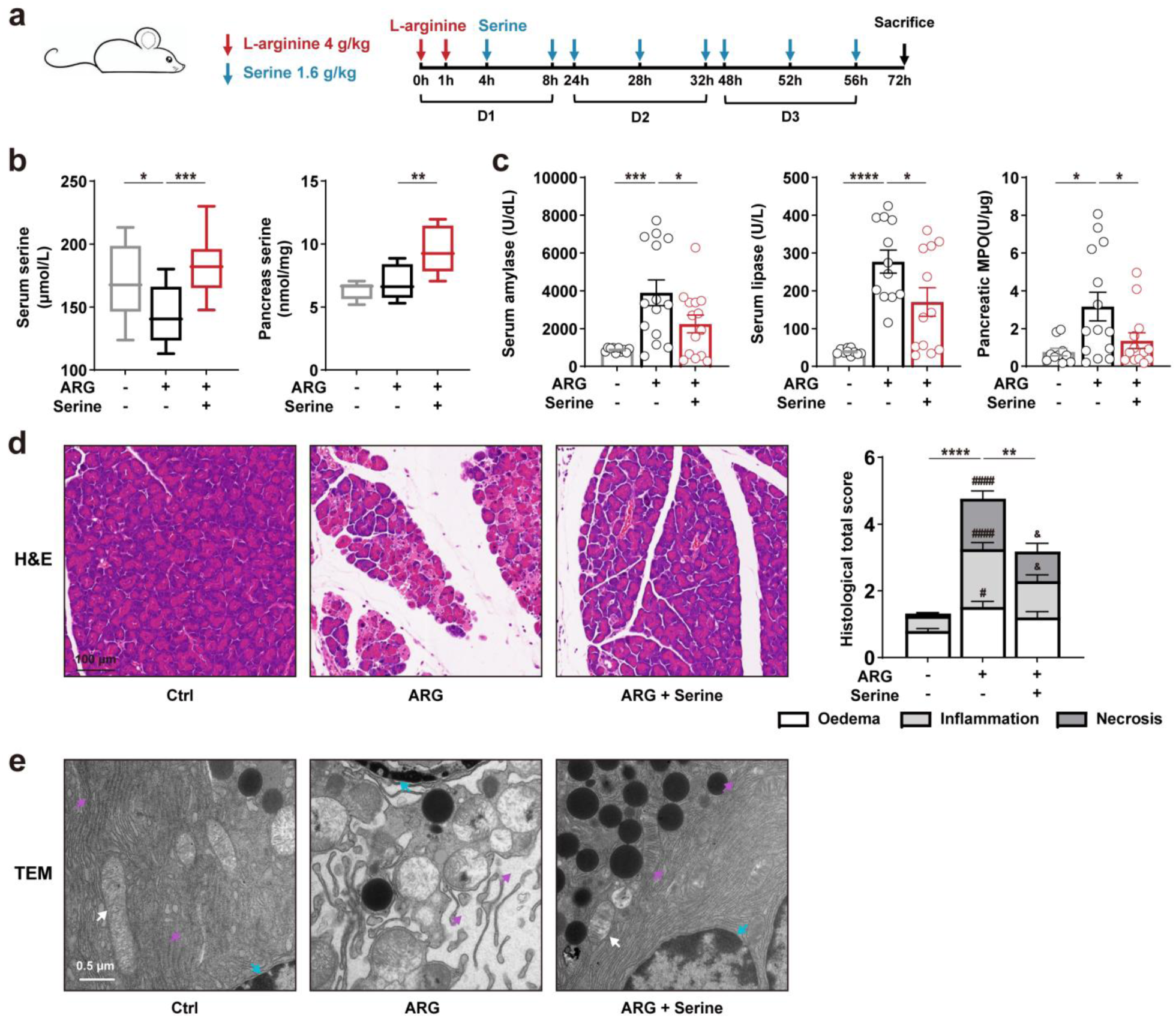
Serine supplementation elevates serum and pancreatic serine levels and ameliorates severity in ARG-AP (related to. Fig. 2**). a,** Schematic illustration of ARG-induced AP modeling and serine administration. **b,** Serum and pancreatic serine concentrations (n = 6–14). **c,** Serum amylase and lipase activities, together with pancreatic MPO activity (n = 10–14). **d,** Representative pancreatic histological images (H&E staining) and quantitative histopathological scoring (right panel) for oedema, inflammation, and necrosis subscores (n = 9–14). Differences in total scores are denoted by *. For subscores, # represents significance compared with the control groups, and & represents significance compared with the AP group. **e,** TEM images showing ultrastructural changes in pancreatic tissue (White arrows, mitochondria; purple arrows, endoplasmic reticulum; green arrows, nucleus). Values shown are mean ± SEM. * *P* < 0.05, ** *P* < 0.01, *** *P* < 0.001, and **** *P* < 0.001.

**Supplementary Fig. 3:**
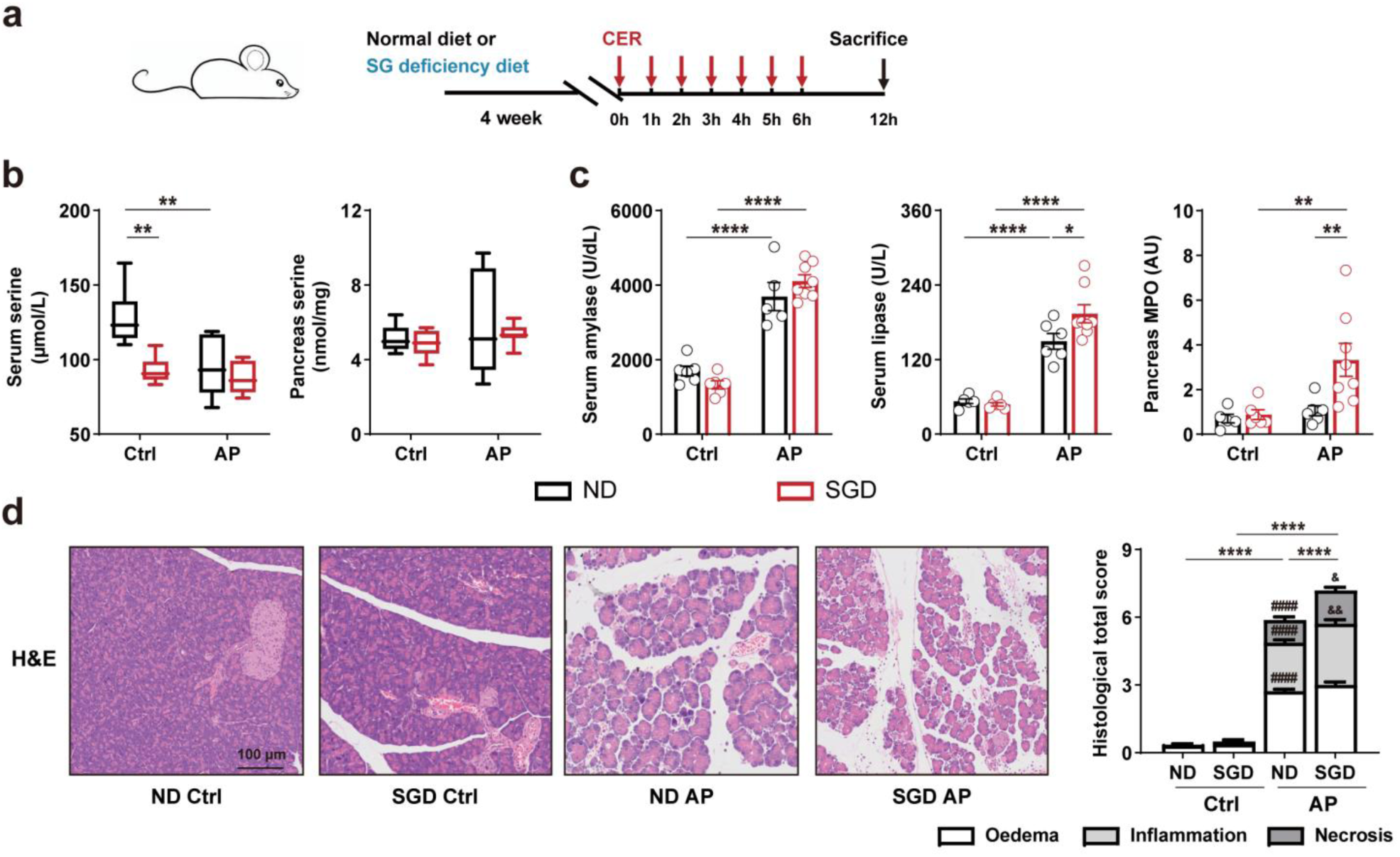
Long-term dietary restriction of serine and glycine lowers serum serine and exacerbates severity in CER-AP (related to. Fig. 2**). a,** Experimental schematic illustrating the combination of CER-induced AP modeling with long-term dietary intervention using an SGD diet. Mice were maintained on normal diet (ND) or SGD diet for 4 weeks prior to CER administration. **b,** Serum and pancreatic serine concentrations (n = 6–10). **c,** Serum amylase and lipase activities, along with pancreatic MPO activity (n = 5–9). **d,** Representative pancreatic histological images (H&E staining) and quantitative histopathological scoring (right panel) for oedema, inflammation, and necrosis subscores (n = 6–10). Differences in total scores are denoted by *. For subscores, # represents significance compared with the ND ctrl group, and & represents significance compared with the ND AP group. Values shown are mean ± SEM. * *P* < 0.05, ** *P* < 0.01 and **** *P* < 0.001.

**Supplementary Fig. 4:**
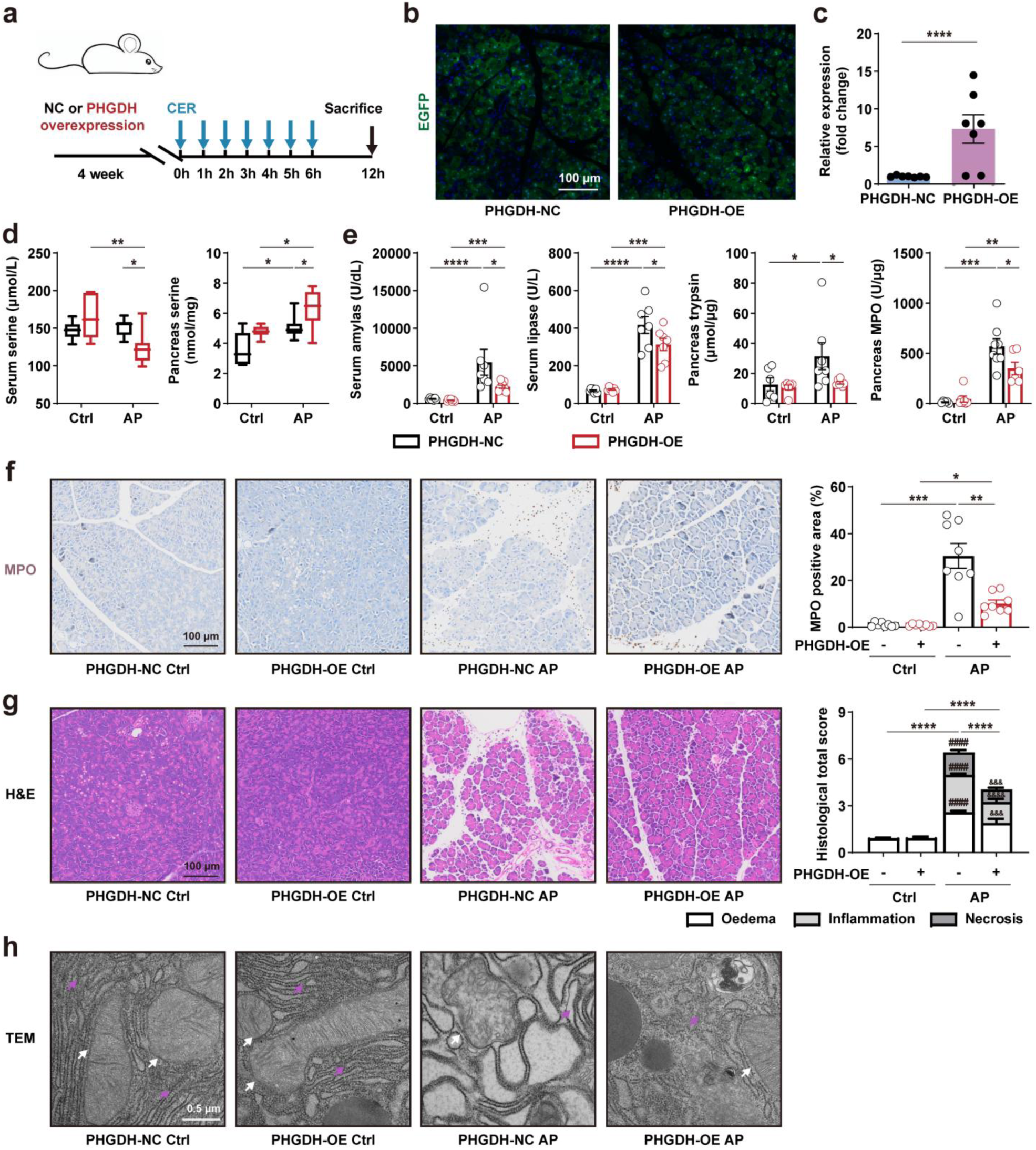
Pancreatic overexpression of PHGDH increases pancreatic serine levels, reduces oxidative stress, and ameliorates CER-AP in mice (related to. Fig. 2**). a,** Experimental design for pancreas-specific PHGDH overexpression (PHGDH-OE) using adeno-associated virus (AAV) delivered via retrograde pancreatic duct injection, followed by CER-induced AP modeling. **b,** Representative fluorescence microscopy showing EGFP expression in AAV-transfected pancreatic tissue. **c,** RT-qPCR quantification of *Phgdh* mRNA expression in pancreatic tissue from PHGDH-OE versus PHGDH-NC (negative control) groups (n = 7). **d,** Serine concentrations in serum and pancreatic tissue, quantified by LC-MS/MS, in PHGDH-OE and PHGDH-NC mice with or without CER induction (n = 6–8). **e,** Serum amylase and lipase activities, as well as pancreatic trypsin and MPO activities (n = 5–8). **f,** Representative IHC images and quantification of pancreatic MPO expression (n=8). **g,** Representative pancreatic histological images (H&E staining) and quantitative histopathological scoring (right panel) for oedema, inflammation, and necrosis subscores (n=6–8). Differences in total scores are denoted by *. For subscores, # represents significance compared with the PHGDH-NC Ctrl group and & represents significance compared with the PHGDH-NC AP group. **h,** TEM images showing ultrastructural changes in pancreatic tissue (White arrows, mitochondria; purple arrows, endoplasmic reticulum). Values shown are mean ± SEM. * *P* < 0.05, ** *P* < 0.01, *** *P* < 0.001, and **** *P* < 0.0001.

**Supplementary Fig. 5:**
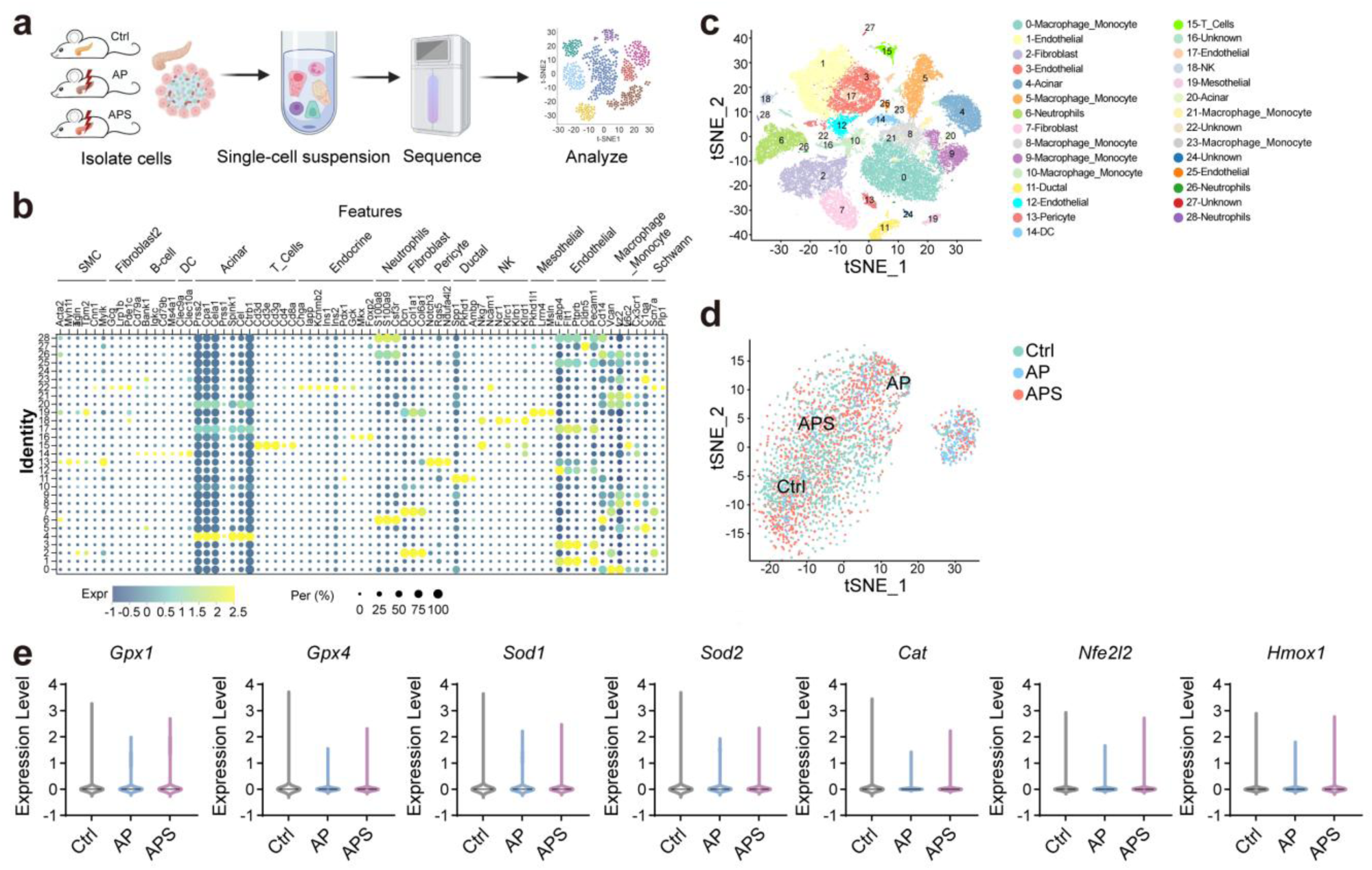
Single-cell transcriptomic analysis and protective effects of serine on primary pancreatic acinar cells (related to. Fig. 3**). a,** Schematic overview of the single-cell RNA sequencing workflow in the control (Ctrl), AP, and APS groups. **b,** Dot plot of canonical marker genes expression across different cell subpopulations. **c,** tSNE plot of all pancreatic cells colored by annotated cell subpopulations. **d,** tSNE plots showing the distribution of three acinar cell clusters in Ctrl, AP, and APS groups. **e,** Violin plots showing expression levels of *Gpx1, Gpx4, Sod1, Sod2, Cat, Nfe2l2 and Hmox1* in Ctrl, AP, and APS groups. Values shown are mean ± SEM.

**Supplementary Fig. 6:**
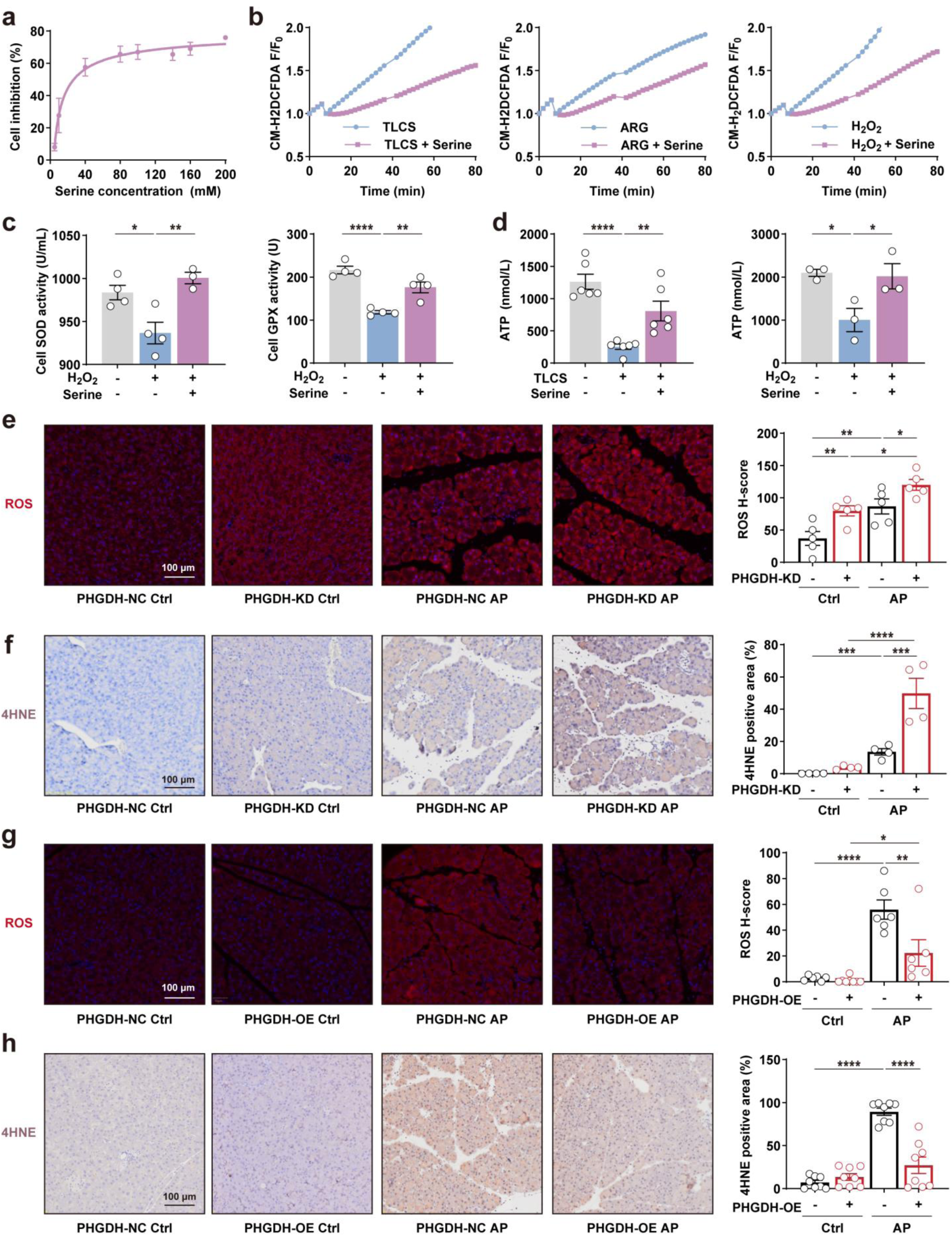
Serine protects acinar cells from oxidative stress damage (related to. Fig. 3**). a,** Dose–response curve of serine treatment on viability of primary PACs (n = 3). **b,** Serine attenuates TLCS-, ARG-and H_2_O_2_-induced ROS accumulation in primary PACs. **c,** Serine enhances total SOD and GPX activities in PACs following H_2_O_2_ treatment (n = 3–4). **d,** Serine alleviates TLCS-and H_2_O_2_-induced ATP depletion in PACs (n = 3–6). **e,** Representative fluorescence images and quantification of pancreatic ROS levels in PHGDH KD mice (n = 5). **f,** Representative IHC images and quantification of pancreatic 4-HNE expression in PHGDH KD mice (n = 4). **g,** Representative fluorescence images and quantification of pancreatic ROS levels in PHGDH OE mice (n = 6–8). **h,** Representative IHC images and quantification of pancreatic 4-HNE expression in PHGDH OE mice (n = 8). Values shown are mean ± SEM. * *P* < 0.05, ** *P* < 0.01, *** *P* < 0.001, and **** *P* < 0.0001.

**Supplementary Fig. 7:**
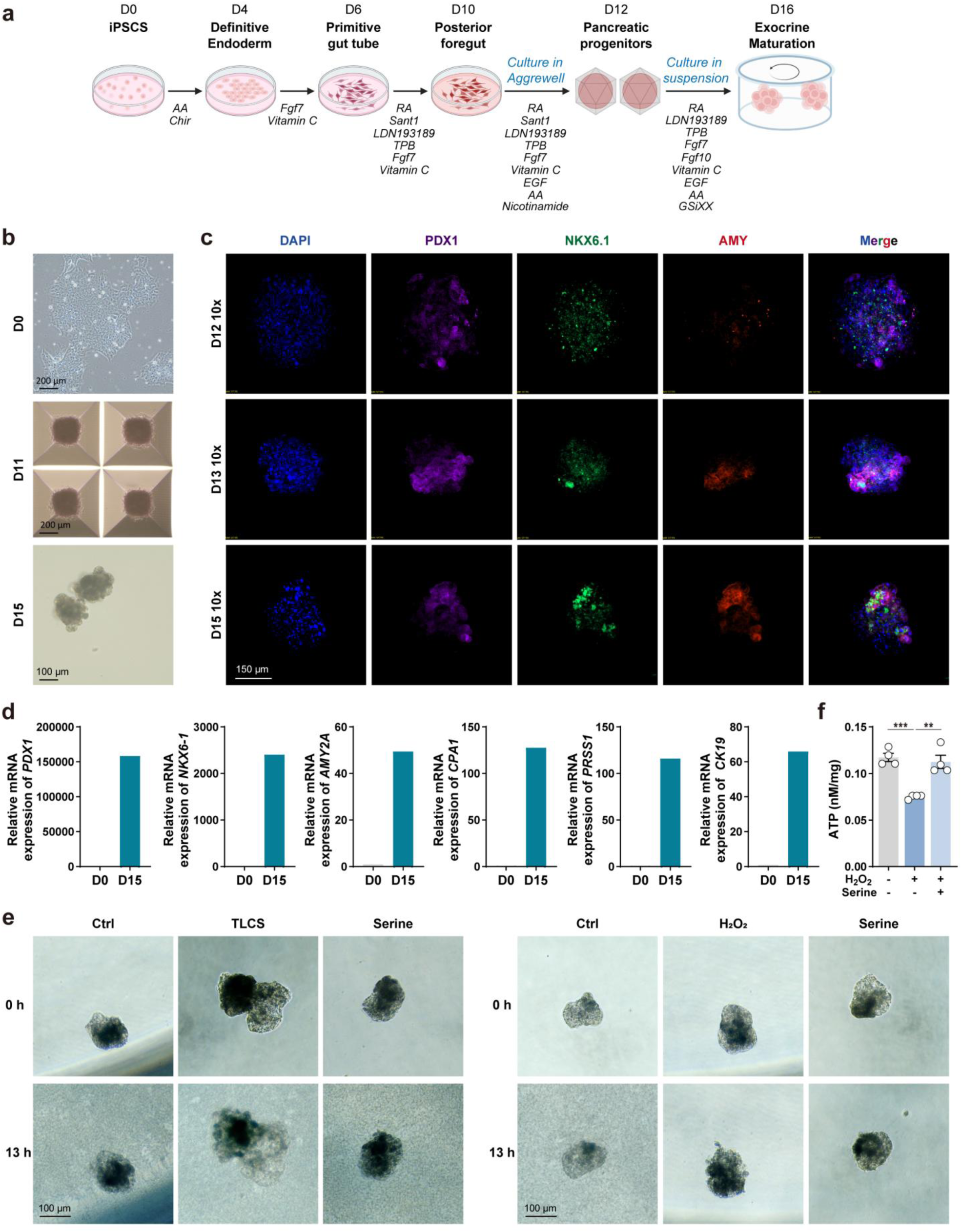
Differentiation of human iPSCs into branched pancreatic organoids (related to. Fig. 3**). a,** Schematic overview of the differentiation protocol and media composition for generating pancreatic organoids from iPSCs. Differentiation was performed in monolayer until day 11 (D11), followed by aggregation in Aggrewell microwells from D12 to D14, and suspension culture until D16. **b,** Representative phase-contrast images of hiPSCs at D0, pancreatic progenitors at D11, and organoids at D15. **c,** Representative confocal images of pancreatic clusters at D12–D15, immunostained for PDX1, NKX6.1, and AMY. Nuclei were counterstained with DAPI. **d,** Representative RT-qPCR analysis of pancreatic marker genes at D0 and D15. Data represent relative fold change normalized to GAPDH. **e,** Morphological changes in TLCS or H_2_O_2_ induced pancreatic organoids following serine treatment. **f,** Serine alleviates H_2_O_2_-induced ATP depletion in D15 pancreatic organoids (n=4). Values shown are mean ± SEM. ** *P* < 0.01, and *** *P* < 0.001.

**Supplementary Fig. 8:**
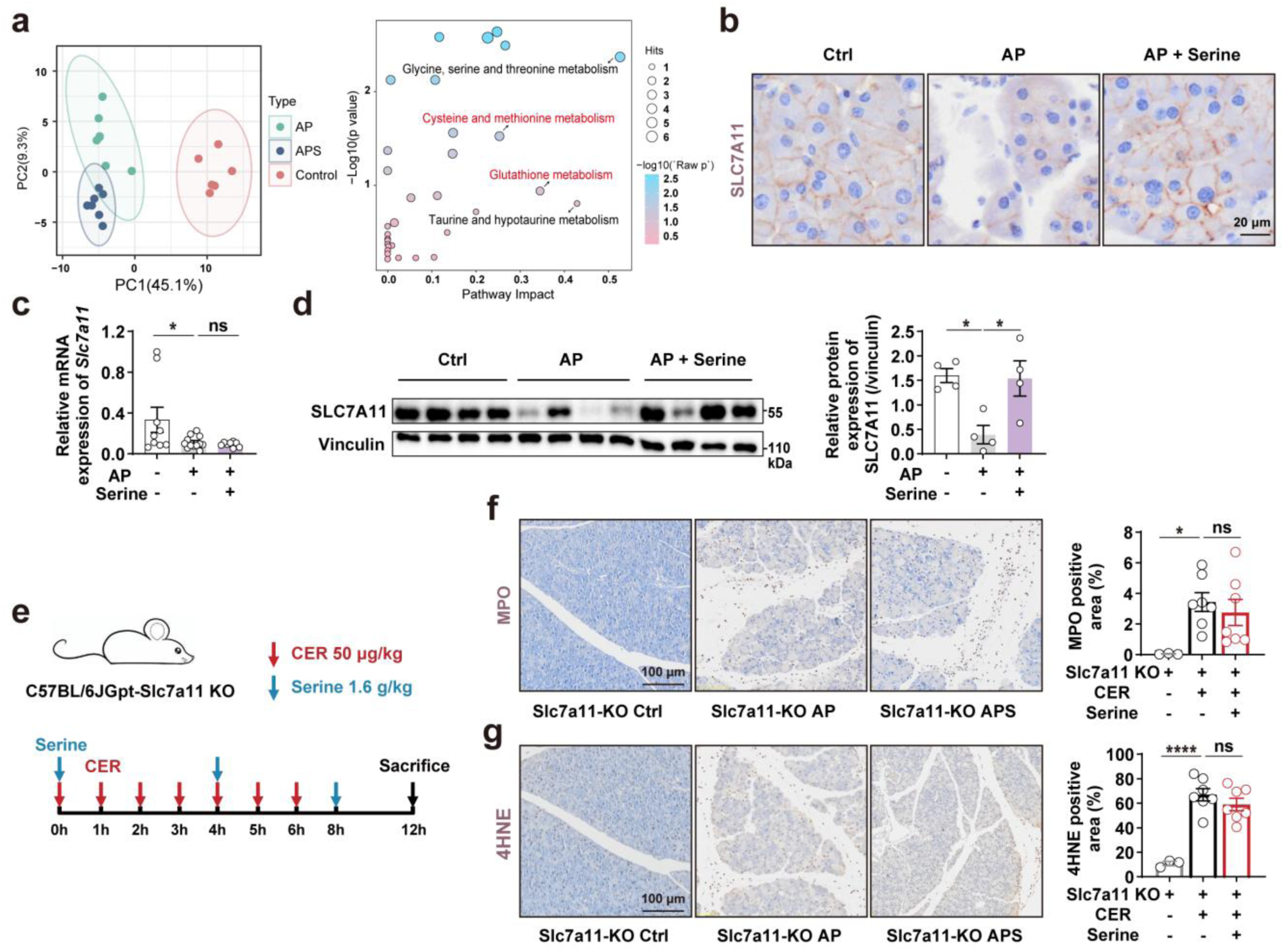
Serine supplementation activates cysteine/methionine and glutathione metabolism pathways and restores SLC7A11 expression *in vivo* (related to. Fig. 4**). a,** Left panel: Principal component analysis (PCA) of targeted metabolomics data from control, AP, and APS groups in the CER-induced AP model. Right panel: KEGG pathway enrichment analysis of differential metabolites between APS and AP groups (adjusted *P* < 0.05). Bubble size indicates metabolite count; color represents-log10 (*P*-value). **b,** Representative IHC images of pancreatic SLC7A11 expression in ARG-AP mice. **c,** Relative *Slc7a11* mRNA expressions in pancreas of ARG-AP mice (n = 8-12). **d,** Western blot analysis and quantification of SLC7A11 protein levels in the pancreas of ARG-AP mice (n = 4). Vinculin served as loading control. **e,** Schematic illustration of CER-induced AP modeling and serine administration in *Slc7a11* KO mice. **f,** Representative IHC images and quantification of pancreatic MPO expression (n = 3-7). **g,** Representative IHC images and quantification of pancreatic 4-HNE expression (n = 3-7). Values shown are mean ± SEM. * *P* < 0.05 and **** *P* < 0.0001. ns meaning *P*>0.05.

**Supplementary Figure 9:**
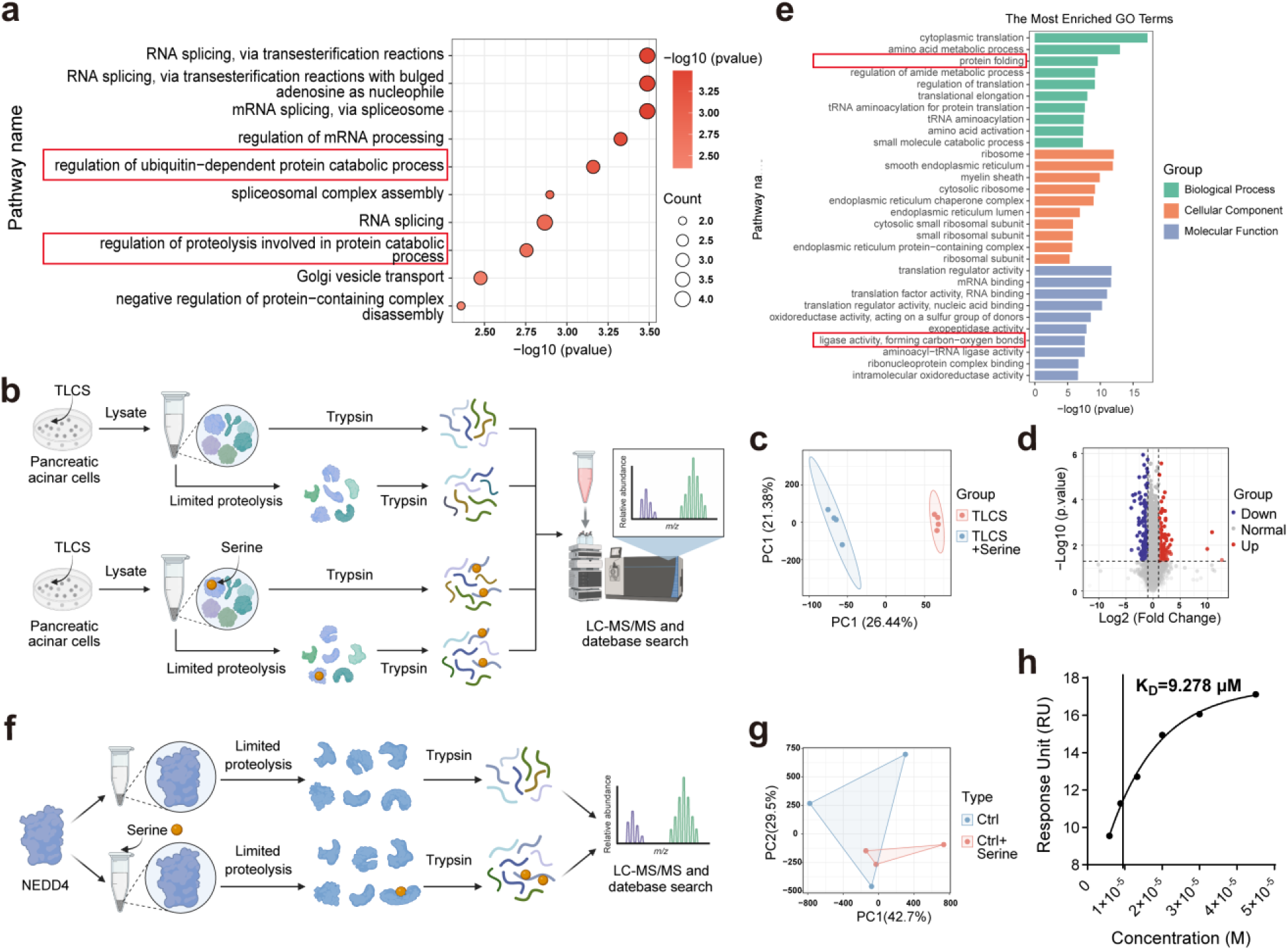
LiP-MS analysis reveals serine-dependent regulation of NEDD4 (related to. Fig. 5**). a,** GO enrichment analysis of differentially expressed proteins in APS vs. AP (CER-AP) from pancreatic proteomics data (n = 4). Bubble size represents the number of proteins (range 2–4), and color indicates-Log10(*P*-value). **b,** Workflow of LiP-MS analysis in TLCS-treated primary PACs with or without of serine (n = 4). **c,** OPLS-DA of LiP-MS data from TLCS-injured primary PACs with or without serine treatment. **d,** Volcano plot showing differentially expressed proteins between TLCS + Serine and TLCS groups. Significantly altered proteins are defined as *P* < 0.05 and FC ≥ 2 or FC ≤0.5. **e,** GO enrichment analysis of LiP-MS–identified differential proteins. **f,** Workflow of LiP-MS analysis performed on human recombinant NEDD4 protein in the presence or absence of serine (n = 3). **g,** PCA of LiP-MS data of human recombinant NEDD4 protein treated with or without serine. **h,** SPR analysis showing the interaction of serine with recombinant NEDD4.

**Supplementary Figure 10:**
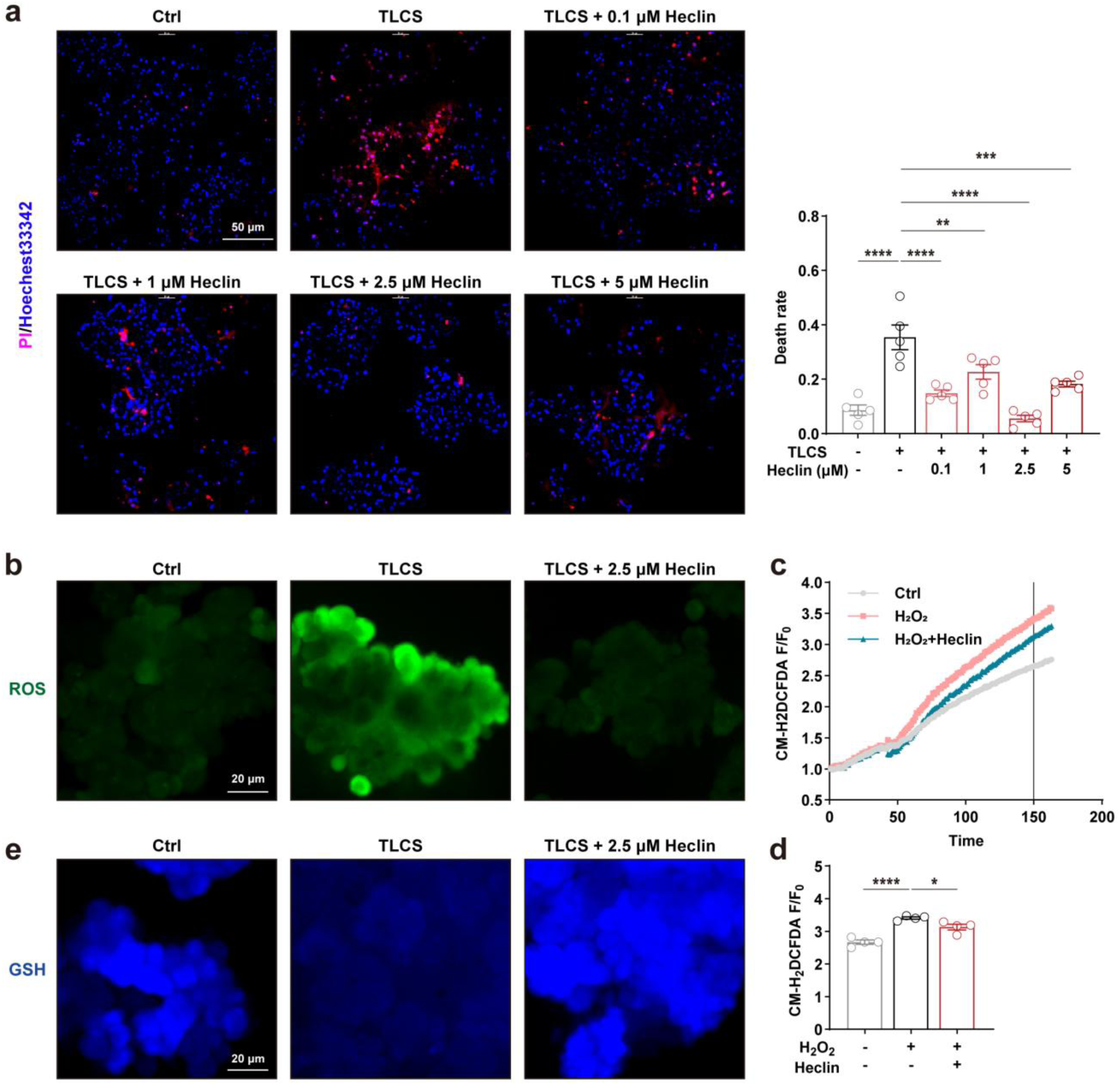
NEDD4 inhibitor heclin directly protects the acinar cells (related to. Fig. 6**). a,** Representative fluorescence images and quantitative analysis of TLCS-induced cell death in primary PACs treated with increasing concentrations of the NEDD4 inhibitor heclin (n = 5). **b,** Representative fluorescence images of ROS. **c,d,** (**c**) Plate reader–based quantification of ROS generation during live-cell imaging of PACs, and (**d**) the corresponding quantitative results at 150 min (n = 4). **e,** Representative fluorescence images showing intracellular GSH production in primary PACs. Values shown are mean ± SEM. * *P* < 0.05, ** *P* < 0.01, *** *P* < 0.001, and **** *P* < 0.0001.

## Notes

### Competing Interest Statement

The authors have declared no competing interest.

